# Single-cell transcriptomics for the 99.9% of species without reference genomes

**DOI:** 10.1101/2021.07.09.450799

**Authors:** Olga Borisovna Botvinnik, Venkata Naga Pranathi Vemuri, N. Tessa Pierce, Phoenix Aja Logan, Saba Nafees, Lekha Karanam, Kyle Joseph Travaglini, Camille Sophie Ezran, Lili Ren, Yanyi Juang, Jianwei Wang, Jianbin Wang, C. Titus Brown

## Abstract

Single-cell RNA-seq (scRNA-seq) is a powerful tool for cell type identification but is not readily applicable to organisms without well-annotated reference genomes. Of the approximately 10 million animal species predicted to exist on Earth, >99.9% do not have any submitted genome assembly. To enable scRNA-seq for the vast majority of animals on the planet, here we introduce the concept of “*k*-mer homology,” combining biochemical synonyms in degenerate protein alphabets with uniform data subsampling via MinHash into a pipeline called `Kmermaid`. Implementing this pipeline enables direct detection of similar cell types across species from transcriptomic data without the need for a reference genome. Underpinning `Kmermaid` is the tool `Orpheum`, a memory-efficient method for extracting high-confidence protein-coding sequences from RNA-seq data. After validating `Kmermaid` using datasets from human and mouse lung, we applied `Kmermaid` to the Chinese horseshoe bat (*Rhinolophus sinicus*), where we propagated cellular compartment labels at high fidelity. Our pipeline provides a high-throughput tool that enables analyses of transcriptomic data across divergent species’ transcriptomes in a genome- and gene annotation-agnostic manner. Thus, the combination of `Kmermaid` and `Orpheum` identifies cell type-specific sequences that may be missing from genome annotations and empowers molecular cellular phenotyping for novel model organisms and species.

## Introduction

Traditional methods to propagate labels from a known single-cell atlas (Regev *et al*., 2017) in one species, to cells in a different species (Shafer, 2019), ignore data from unaligned reads, unannotated genes, and incomplete ortholog annotations (Figure 1, top). Despite recent advances in end-to-end genome assembly (Formenti *et al*., 2020; Miga *et al*., 2020; Rhie *et al*., 2020), RNA-seq data is more readily available than high quality *de novo* genome assemblies; however, traditional transcriptomic analyses rely on alignment to a genome, thus prohibiting its utility for comparative analyses. Even when a species has an available reference genome, three main problems arise in comparing transcriptomes across species. First, most animals in the wild are more heterozygous than laboratory-grown animals (da Fonseca *et al*., 2016); thus their reads tend to have a lower rate of alignment due to high variation between the reference genome and individuals, thus reducing the amount of data used in downstream analyses. Second, gene annotations are often incomplete due to differences in species’ gene structures, and partial sampling of all possible expressed genes across developmental stages and cell types. Third, in most analyses, only genes pre-identified from a database such as ENSEMBL (Yates *et al*., 2020) with a one-to-one match (“1:1 orthologs”) across species are considered, disregarding species-specific genes that may be of interest (Shafer, 2019). Even with improvements in the quality of genome assembly and gene annotation, the open problem of identifying orthologs across species remains challenging (Sonnhammer *et al*., 2014; Altenhoff *et al*., 2016, 2020; Nichio, Marchaukoski and Raittz, 2017), especially across large evolutionary distances. Orthology detection tools (Sonnhammer *et al*., 2014; Emms and Kelly, 2015, 2019; Altenhoff *et al*., 2016; Huerta-Cepas *et al*., 2016, 2017) are useful but computationally slow and again require genome assemblies and gene annotations --missing for the vast majority of animals. With the exception of SAMap (Tarashansky *et al*., 2021), all cross-species label propagation tools require orthologous gene assignment, and even SAMap is limited to annotated reference genomes. Thus, strategies outside of the traditional alignment-based methods are needed to identify common cell types across species.

**Figure 1.**
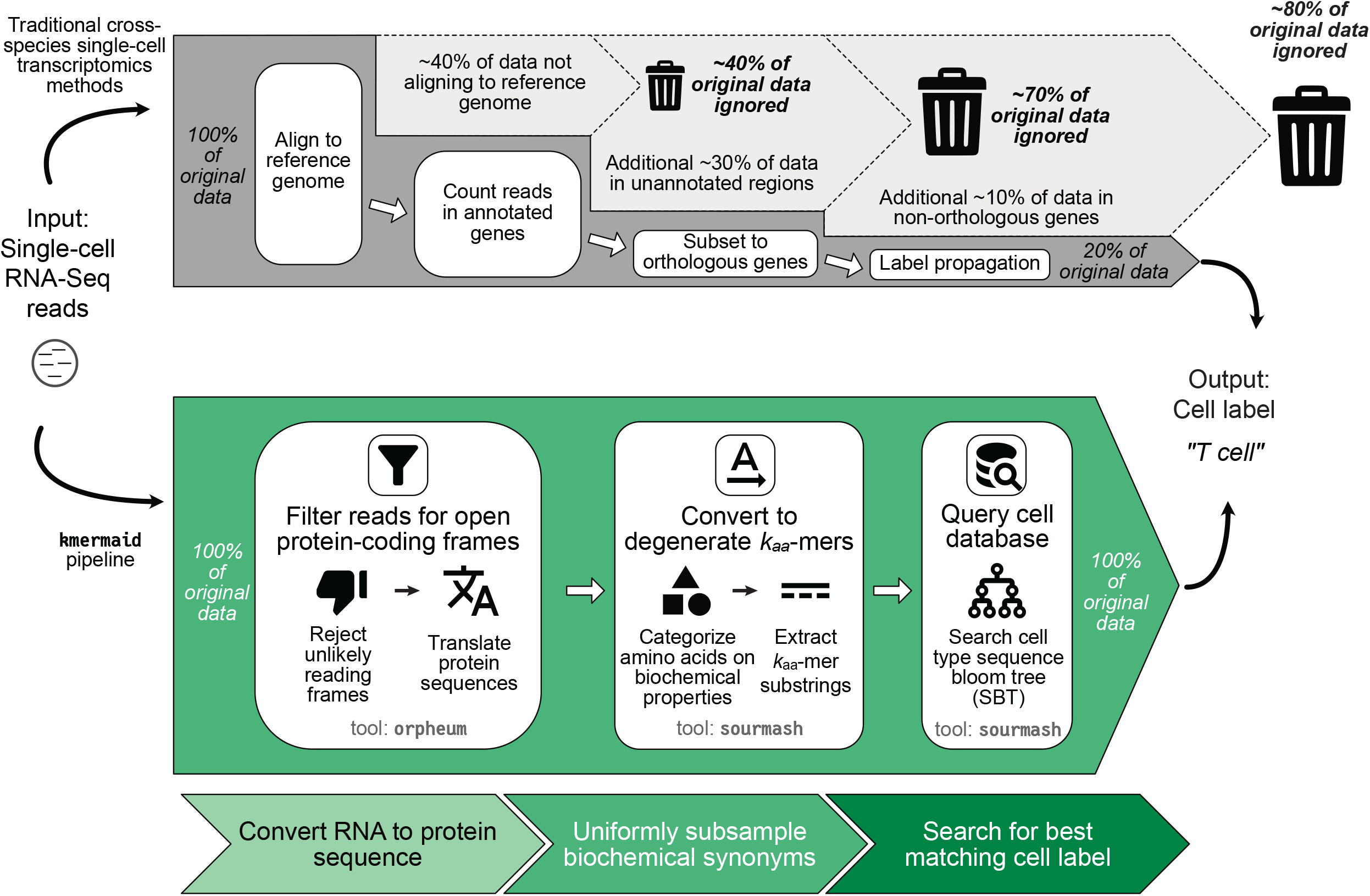
Kmermaid is an alignment-free method for cross-species cell type identification. Top, overview of traditional methods for cross-species cell type identification. Bottom, Overview of the kmermaid pipeline. First, RNA-seq reads are translated to protein via the orpheum tool, then the translated protein sequences are re-encoded into a reduced alphabet representing their biochemical properties and short words are subsampled to create a *k*-mer signature. The cells’ *k*-mer signature is then queried against a curated cell type database and the best matching cell label is returned.

Cell types can be aggregated by their broad functional categories or cell lineage, such as grouping B cells and T cells into a common lymphoid *compartment*. In this work, we refer to five cellular compartments: endothelial, epithelial, lymphoid, myeloid, and stromal.

Short subsequences of length *k*, or *k*-mers are widely used in bioinformatics (Compeau and Pevzner, 2018) to alleviate data loss issues in comparing aligned reads (Ondov *et al*., 2016). Using nucleotide *k*-mers (hereafter *k*_*nuc*_-mers) for identifying cell types has been shown to perform as well as using read counts of genes from single cell sequencing data (Shi and Yip, 2019). Underlying `kmermaid` is the recent application of *k*-mer subsampling techniques to compress data into a smaller form factor, but faithfully represent the underlying data as if all *k*-mers were present (Brown and Irber, 2016; Ondov *et al*., 2016; Pierce *et al*., 2019; Irber and Brown, 2020). As protein sequences are more evolutionarily conserved than the underlying DNA, in this work, we developed the tool `orpheum` to extract putative translated amino acid sequences. We then uniformly downsampled amino acid sequence *k*-mers (Ondov *et al*., 2019) (hereafter *k*_*aa*_-mers) from each single cells’ translated protein sequence using a modified MinHash algorithm as implemented in Sourmash (Brown and Irber, 2016; Pierce *et al*., 2019). As orthologs share amino acid sequence, they also share *k*_*aa*_-mers, and each *k*_*aa*_-mer represents ortholog fragments, bypassing the computationally intensive step of all-by-all matching of entire orthologous sequences present in most orthology detection tools.

Evolutionarily divergent, but functionally conserved, protein orthologs often have amino acid changes that nonetheless retain their biochemical properties. To account for amino acid differences in protein sequence across species and to expand the evolutionary range of our method, we employed degenerate amino acid alphabets. Reduced amino acid alphabets are useful in rapid database searches of related proteins (Landès and Risler, 1994; Edgar, 2004; Ye, Choi and Tang, 2011; Buchfink, Xie and Huson, 2015), protein structure prediction (Murphy, Wallqvist and Levy, 2000; Peris, López and Campos, 2008; Peterson *et al*., 2009), and as a computationally lightweight method of identifying orthologous genes (Hu and Friedberg, 2019). Specifically, we used the Dayhoff encoding (Dayhoff and National Biomedical Research Foundation, 1969; Dayhoff, 1972; DAYHOFF and M. O, 1972) which categorizes amino acids into six chemical groups: 1) sulfur polymerization 2) small, 3) acid or amide, 4) basic, 5) hydrophobic, and 6) aromatic (Table 1).

**Table 1:**
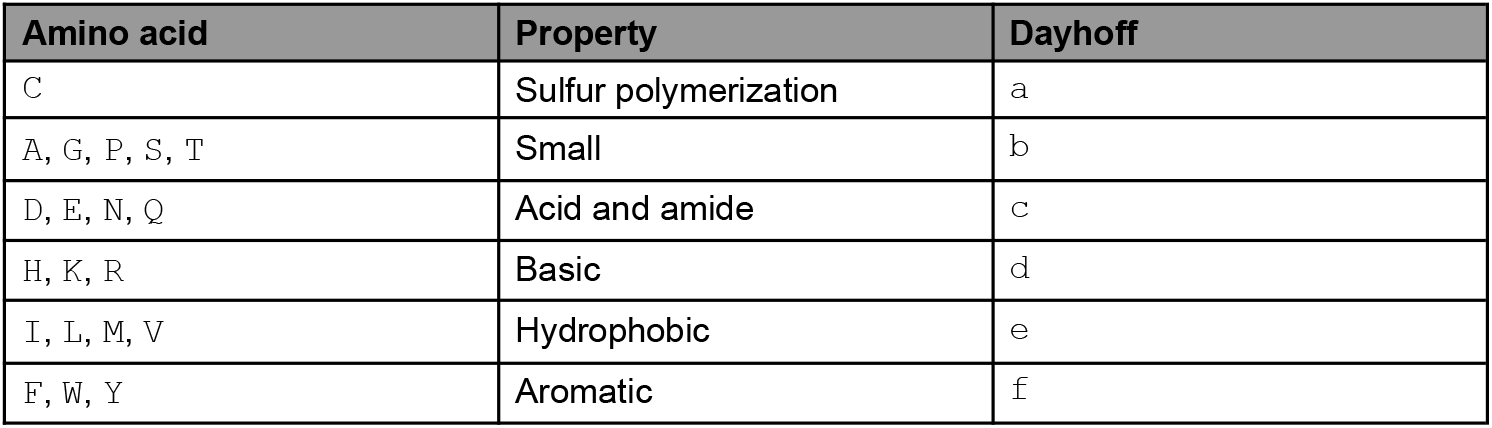
The Dayhoff encoding is a reduced amino acid alphabet allowing for permissive cross-species sequence comparisons. For example, the amino acid sequence `LIVING` would be Dayhoff-encoded to `eeeecb`.

We introduce the `kmermaid` pipeline to extract evolutionarily conserved sequences from single-cell RNA-seq data and predict the cell type. `kmermaid` enables the identification of putative homologous genes from short *k*-long amino acid sequence fragments, which we term “*k*-mer homology.” `kmermaid` addresses the unmet need to quantitatively compare single-cell transcriptomes across species without the need for orthologous gene mapping, gene annotations, or a reference genome. Here we have implemented *k*_*aa*_-mers with reduced amino acid alphabets computationally translated with our tool, `orpheum`, to find shared cellular compartments across evolutionarily divergent mammalian species, with divergence times of ≥90 million years. As the direct assignment of protein-coding sequence skips both traditional alignment and gene orthology assignment, it can a) be applied to transcriptomes from organisms with no or poorly annotated genomes, and b) identify putative functions of protein sequences contributing to shared cell types. To demonstrate the utility of these alignment-free methods, we identified cell types in the lung of the Chinese horseshoe bat, *R. sinicus*, the known host of SARS-CoV-1, and the purported host of SARS-CoV-2.

## Results

To circumvent the need for a reference genome in conventional scRNA-seq analyses, we developed a computational pipeline called Kmermaid (Figure 1, bottom) to compare divergent protein sequences translated from RNA-seq data to identify annotate cell types.

### Reliable identification of putative protein-coding sequence across ~100 million years of evolution with orpheum translate

The first step in kmermaid requires accurate protein-coding frames from scRNA-seq data (Figure 2). We posited that for species without a reference genome, it may be sufficient to use a database of protein sequences from organisms related within ~100 million years. As protein orthologs share sequences, they also share *k*_*aa*_-mers. For each RNA read, there are six possible reading frames: three for each of the forward and backward frames. However, ~50% of the frames contain a stop codon, and many don’t resemble known proteins. To retain only likely reading frames, we created the tool orpheum to extract likely protein-coding sequences from each RNA read. Orpheum performs six-frame translation of all reads and discards reading frames with stop codons or too many “nullomers” (Hampikian and Andersen, 2006), i.e. *k*_*aa*_-mers not present in the protein sequence database (Fig. 2A). After translation, (Fig. 2A, step 1) amino acid sequences are decomposed into amino acid *k*_*aa*_-mers (step 2), which are then compared with reference proteome sequences (step 3). We benchmarked orpheum on simulated human RNA-seq data to find the most performant k_aa_-mer sizes using a strict dataset: the intersection of single-copy orthologous genes present in all extant mammals as defined by BUSCO (Simão *et al*., 2015) and the Quest for Orthologs 2019 datasets (Fig 2B, Supplemental Fig. 1A, Methods). Classification power, measured by receiver operating characteristic area under the curve (ROC AUC) (Hastie, Tibshirani and Friedman, 2009) achieved a maximum across most species with *k*_*aa*_=8 for the Protein alphabet and *k*_*aa*_=17 for the Dayhoff alphabet (Fig. 2C, Supplemental Fig. 1B-C). Even with the most diverged group at ~96 mya, both alphabets performed well with a maximum ROC AUC 0.929 with the Protein alphabet and 0.860 with the Dayhoff alphabet. For downstream analyses, we proceeded with the Protein alphabet and k=8 as our `orpheum translate` parameters. The maximum ROC AUC was more highly correlated to the number of proteins and the number of unique k-mers in the proteome than the divergence time (Supplemental Fig. 1D-F), indicating that proteome database complexity is more important than using a closely related organism, e.g. related within ~50 mya. Using a threshold of 5% *k*_*aa*_-mer presence in the proteome database, orpheum precisely removed noncoding sequences (Fig 2E,F, left columns) across all divergence times, while recall of protein-coding sequences declined with divergence time (Fig 2E,F, right columns). Thus, protein-coding sequences were reliably identified even if the proteome database was from a species up to ~100 million years of divergence, provided it contained at least ~4000 sequences (Supplemental Fig. 1G). These results establish the ability of Orpheum to accurately detect putative protein-coding sequences and reliably reject non-coding sequences, even when the proteome database is approximately 100 million years diverged from the query organism.

**Figure 2.**
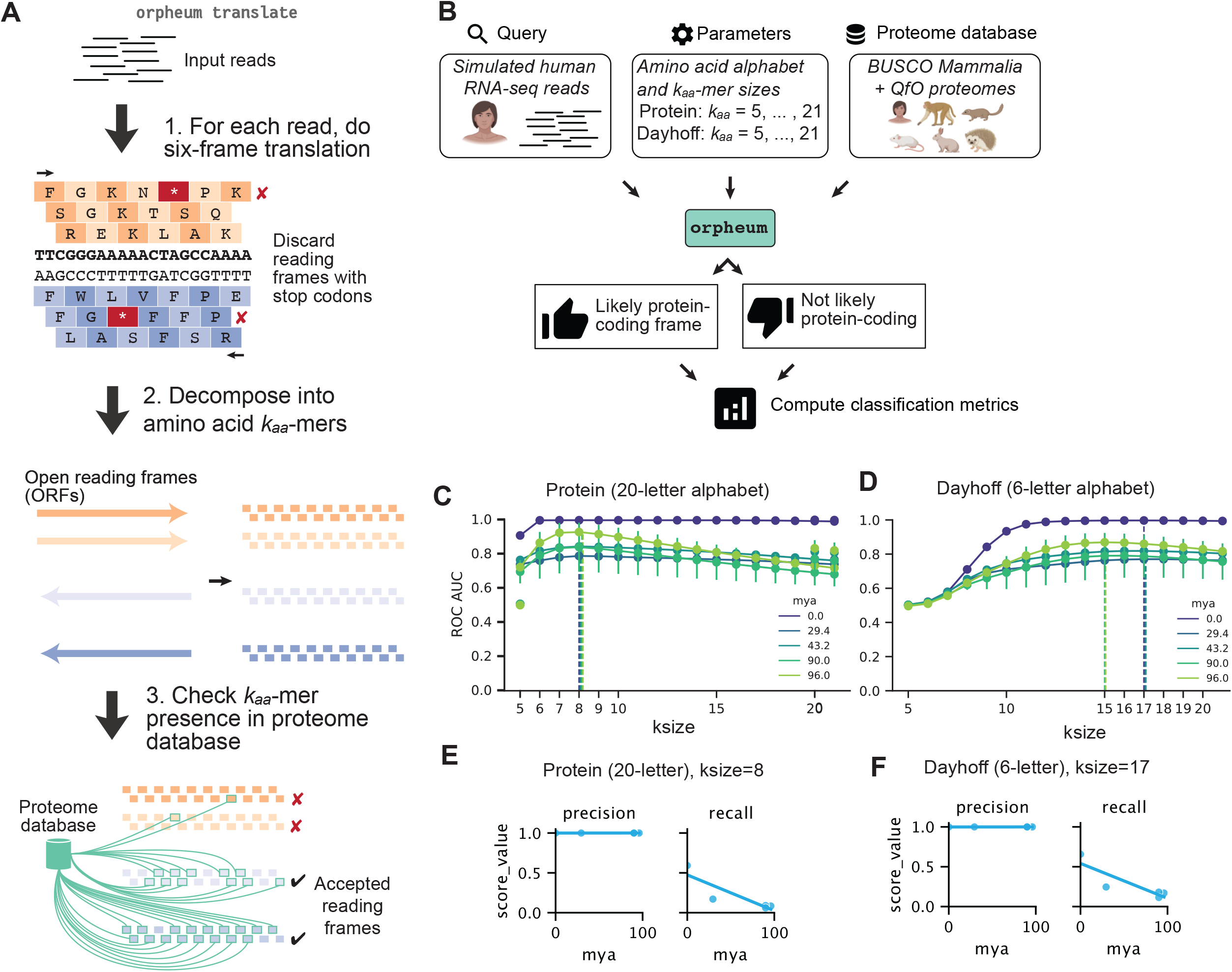
Orpheum precisely predicts protein-coding reading frame. **A**. Overview of `orpheum translate` method. First, each read is translated into all six possible protein-coding translation frames. Next, reading frames with stop codons are eliminated. Each protein-coding frame is decomposed into *k*_*aa*_-mer amino acid words, then the fraction of *k*-mers appearing in the proteome database are computed. Reading frames containing 5% or more *k*-mers matching the reference proteome are inferred to be putatively protein-coding. **B**. Experimental set up to identify best *k*-mer sizes for translating protein-coding sequence, given a reference proteome of divergent organisms. First, RNA-seq reads were simulated from human genes whose orthologs are present in all extant mammals, as defined by BUSCO Mammalia. Next, we used BUSCO genes overlapping the highly curated reference proteomes from the Quest for Orthologs (QfO), representing almost 200 million years of evolution, across a wide range of mammalian species. Using `orpheum`, we partition the reads into protein-coding and noncoding using a variety of amino acid alphabets and *k*-mer sizes, and then compute classification metrics. Animal images from BioRender.com **C**. Receiver operating characteristic area under the curve (ROC AUC) Classification metrics of protein-coding prediction with the 20-letter Protein alphabet. x-axis, *k*_*aa*_-mer size used for orpheum translate; y-axis, ROC AUC value; hue, millions of years (mya) of estimated divergence time of reference proteome relative to human, from timetree.org. Dotted lines highlight the maximum ROC AUC for each mya at ksize=8 for all divergence times. **D**. Receiver operating characteristic area under the curve (ROC AUC) Classification metrics of protein-coding prediction with the 6-letter Dayhoff alphabet. x-axis, *k*_*aa*_-mer size used for orpheum translate; y-axis, ROC AUC value; hue, millions of years (mya) of estimated divergence time of reference proteome relative to human, from timetree.org. Dotted lines highlight the maximum ROC AUC for each mya at ksize=17 for most divergence times. **E**. Precision and recall of known protein-coding sequences using a ksize=8 with the 20-letter Protein alphabet. Left, precision and right, recall. x-axis, millions of years (mya) of estimated divergence time of reference proteome relative to human, from timetree.org; y-axis, value of precision or recall. **F**. Precision and recall of known protein-coding sequences using a ksize=17 with the 6-letter Dayhoff alphabet. Left, precision and right, recall. x-axis, millions of years (mya) of estimated divergence time of reference proteome relative to human, from timetree.org; y-axis, value of precision or recall.

### Kmermaid uses *k*_*aa*_-mers from reduced amino acid alphabets to enable cell type label propagation across species

After prediction of protein-coding frames using Orpheum, we converted amino acids into Dayhoff categories (Table 1) and performed MinHashing of *k*-mers using Sourmash (Brown and Irber, 2016; Pierce *et al*., 2019) to create *k*-mer signatures, subsampled to approximately 1/10th of the original *k*-mers. To ensure that our *k*_*aa*_-mer alphabet encoded sufficient sequence complexity to be discriminating, we first determined that *k*_*nuc*_-mers ≥ 21, using a four-letter DNA alphabet (let |∑| alphabet size, then |∑_nuc, DNA_| = 4), are sufficient to uniquely identify DNA sequences. We then computed the total space of possible *k*_nuc_-mers, which we termed “complexity,” |∑|^k^ = |∑_nuc, DNA_|^k_DNA^ = 4^21^. To prevent numeric overflow, we used the log of the total number of *k*-mers, or *k* log(|∑|) = *k*_nuc_ log (|∑_nuc, DNA_|) (Supplemental Figure 2A). The 20-letter Protein alphabet (|∑_aa, Protein_| = 20) has an equivalent complexity at *k*_*aa*_=10, and the six-letter Dayhoff alphabet (|∑_aa, Dayhoff_| = 6) has an equivalent complexity at *k*_*aa*_ = 17. Thus, we chose *k*_*aa*_=10 for the Protein alphabet and *k*_*aa*_ = 17 for the Dayhoff alphabet. As *k*_*aa*_-mer signatures enable fast comparison of transcriptional profiles using only a subsampling of the total sequence content for each cell, the flexibility of reduced amino acid alphabet k_aa_-mers enables the identification of orthologous protein fragments across divergent species. Finally, similar to traditional methods based on read counts in orthologous genes, these *k*_*aa*_-mer signatures can be used to identify common cell types.

As a testbed, we chose the lung as it is a complex organ with many cell types, and well-annotated lung cell atlases exist for humans (Travaglini *et al*., 2019), mice (Neff *et al*., 2018; Tabula Muris Consortium, 2020) and the Chinese horseshoe bat (Ren *et al*., 2020) (*Rhinolophus sinicus*), which diverged from a common ancestor between 85 and 97 million years ago (*TimeTree* :: *The timescale of life*, no date a) (Figure 3A). To determine the utility of *k*-mers for cell type identification, we first tested `kmermaid` with a broadly categorized cell type database (e.g. by grouping all T cell subtypes into “T cell” etc. Table 2), and strengthened cell-type specific signals by removing uninformative *k*-mers (Supplemental Figure 2B, Methods). Using a cell type database of mouse lung, we queried individual mouse, human, and bat lung cells. Classification performance was assessed using Adjusted Rand Index (ARI) (Hastie, Tibshirani and Friedman, 2009), using bootstrapping to randomly subsample the cell type labels 1000 times and compute classification metrics (Fig. 3B). The mouse-to-mouse cell type performance was steady across the three alphabets with mean ARIs ~0.8, while the cross-species cell type label propagation did not perform as well, with mean ARI ~0.3-0.4. To further explore these discrepancies, we generated confusion matrices (Supplemental Fig. 3A-I), which revealed that for the more accurate Dayhoff alphabet, the most common mis-classifications tended to swap labels within cellular compartment, e.g. Macrophages labeled as their myeloid lineage precursor, Monocytes (Supplemental Fig 3. F, I). Thus, we also computed the ARI and visualized Sankey diagrams (Figure 3C-F) and confusion matrices (Supplemental Figure 4) for retention of cellular compartments. Interestingly, while the DNA and Protein alphabets performed poorly for the cross-species propagation, the Dayhoff alphabet did not have as substantial a drop in performance, remaining above 0.8 ARI for all species, almost a two-fold improvement over more granular cell type classification. The higher performance of the Dayhoff alphabet in humans and bats was also consistent with the F1 score metric (Hastie, Tibshirani and Friedman, 2009) (Supplemental Fig 5). Thus, short sequences from a degenerate amino acid alphabet are more useful at identifying cellular compartments across species than DNA or primary protein sequences.

**Figure 3.**
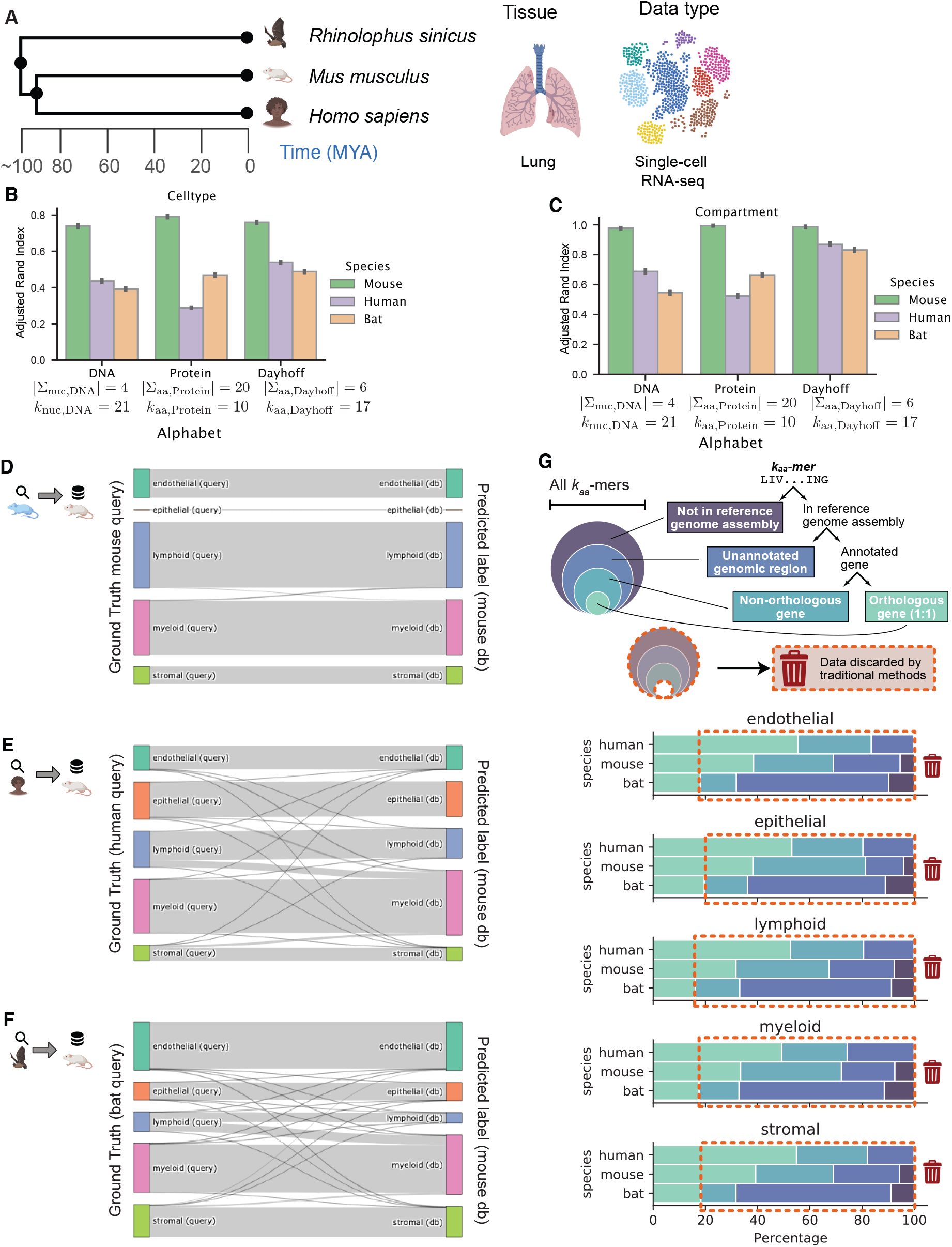
*k*_*aa*_-mers generated by kmermaid identify cell types across species. **A**. Overview of single-cell RNA-seq lung tissue datasets from three species: Homo sapiens (human), Rhinolophus sinicus (Chinese horseshoe bat), and Mus musculus. Animal, Lung, and single-cell embedding images from BioRender.com **B**. Adjusted rand index (ARI) of kmermaid cell type label propagation within species and across species, using the DNA (|∑_nuc, DNA_| = 4, *k*_*DNA*_ = 21), Protein (|∑_aa, Protein_, DNA| = 20, *k*_*Protein*_ = 10), and Dayhoff (|∑_aa, Dayhoff_| = 6, *k*_*Dayhoff*_ = 17) alphabets. **C**. Adjusted rand index (ARI) of kmermaid cellular compartment label propagation within species and across species, using the DNA (|∑_nuc, DNA_| = 4, *k*_*DNA*_ = 21), Protein (|∑_aa, Protein_, DNA| = 20, *k*_*Protein*_ = 10), and Dayhoff (|∑_aa, Dayhoff_| = 6, *k*_*Dayhoff*_ = 17) alphabets. **D**. Mouse reference to mouse query cellular compartment label propagation via kmermaid Sankey diagram, using the Dayhoff alphabet (|∑_aa, Dayhoff_| = 6, *k*_*Dayhoff*_ = 17). **E**. Mouse reference to human query cellular compartment label propagation via kmermaid Sankey diagram, using the Dayhoff alphabet (|∑_aa, Dayhoff_| = 6, *k*_*Dayhoff*_ = 17). **F**. Mouse reference to bat query cell cellular compartment propagation via kmermaid Sankey diagram, using the Dayhoff alphabet (|∑_aa, Dayhoff_| = 6, *k*_*Dayhoff*_ = 17). **G**. Presence of Dayhoff (|∑_aa, Dayhoff_| = 6, *k*_*Dayhoff*_ = 17) *k*_*aa*_-mers not in the reference genome assembly, in the reference assembly but not in an annotated gene, in an annotated gene but not a known ortholog, or a known ortholog. Top, flow chart of diagnostic k-mer categories. Middle, red dotted line and shaded region showing data not in reference assembly, in unannotated genome regions, and in non-orthologous genes discarded by traditional methods. Bottom, stacked bar plots of the percentage (x-axis) of diagnostic k-mers per predicted cell type label, in each species (y-axis). Red dotted box indicates data that would be thrown out using traditional comparative methods.

### Kmermaid enables identification of unannotated cell type-enriched sequences in the Chinese horseshoe bat

Given that Dayhoff *k*_*aa*_-mers are a more degenerate representation of sequence, we wondered whether they could be used to identify unannotated transcripts in one species from an annotated transcript in cell types across species (Supplementary Figure 6A). To test this, we identified Dayhoff *k*_*aa*_-mers shared by both bats and humans, aligned in the human assembly, but not in the bat assembly (Supplementary Figure 6B-E). Table 3 shows the number of bat cells expressing *k*_*aa*_-mers unaligned in the bat genome, but were shared with human genes. Of the ground truth cell type annotations, Alveolar Epithelial Type 2 (AT2) cells expressed the most antiviral and interferon-stimulated genes (ISG) detected by bat-unaligned k-mers, but matched aligned k-mers in human (Supplementary Figure 6B) including the ISG proteases *CASP4* detected by 209 cells and *CAPN2* detected in 133 cells in bats, suggesting an interferon activation signature in the AT2 cells. Additionally, 161 capillary cells in bat included *k*_*aa*_-mers harbored in reads aligning to *TEAD4* in humans, whose nuclear localization is necessary for VEGF-induced angiogenesis (Liu *et al*., 2011), and thus is a known cell type marker gene for capillary cells. In the bat genome, components of the major histocompatibility complex (MHC), expressed in professional antigen presenting cells such as monocytes, were not annotated, but *k*_*aa*_-mers in unaligned bat sequences matched to *HLA-DRB1* and *HLA-DRB6* genes in human were expressed in 41 and 36 monocyte cells, respectively. None of the genes *CASP4, CAPN2, TEAD4, HLA-DRB1, HLA-DRB6* were annotated in the bat genome, and yet we were able to identify *k*_*aa*_-mers from these human genes, that were present in the bat transcriptome but did not successfully align to the bat genome. These results show that unaligned sequences contain useful information. Thus, unaligned *k*_*aa*_-mers can identify known cell type marker genes.

As traditional methods for cross-species analyses utilize one-to-one orthologs, we were interested in the percentage of orpheum-translated Dayhoff *k*_*aa*_-mers present in either the genome assembly, in an annotated gene, or in a 1:1 ortholog, for each species (Fig. 3G). For each single cell’s predicted cellular compartment, we assigned *k*_*aa*_-mers to a gene if it was contained in a read aligning to that gene, and assigned orthology using Mouse Genome Informatics JAX Lab (*MGI-Mouse Genome Informatics-The international database resource for the laboratory mouse*, no date) and ENSEMBL homology databases (Vilella *et al*., 2008). Consistent with the annotation depth, less than 1% of *k*_*aa*_-mers from human were unaligned relative to the human genome, whereas mouse had 4-8% of *k*_*aa*_-mers not present in the mouse genome assembly, and 8-12% of bat *k*_*aa*_-mers were not present in the *R. sinicus* genome. Overall, 20% or fewer *k*_*aa*_-mers appeared in annotated 1:1 orthologs in the bat, indicating that only 20% of the *k*_*aa*_-mers are usable for traditional cross-species analyses. Thus, the use of *k*_*aa*_-mers enabled the utilization of ~80% data that would traditionally be discarded as it is not present in the genome assembly, in unannotated genes, or genes that are not 1:1 orthologs.

## Discussion

There is an urgent need for methods that propagate labels from model organism cell atlases to understudied animals as they empower researchers to use unconventional species with a striking phenotype to model human disease. In this paper, we present methods to map cell types from well-annotated organisms to any other organism, regardless of the need for a reference genome. The availability of whole organism cell atlases (Regev *et al*., 2017; Neff *et al*., 2018; Tabula Muris Consortium, 2020) enables rapid cell type identification from new datasets. However, cell type identification from scRNA-seq data is not readily accessible to the 99.9% of the planet’s animal species (Mora *et al*., 2011; Scheffers *et al*., 2012) that lack a reference genome. Moreover, a reference genome alone is not sufficient for multi-species transcriptomic analyses, as computationally expensive mapping of equivalent (orthologous) genes across species is also required (Sonnhammer *et al*., 2014; Altenhoff *et al*., 2016, 2020; Nichio, Marchaukoski and Raittz, 2017; Shafer, 2019). Thus, despite the relative ease of obtaining transcriptomic data, its application to understudied organisms, without a genome, is rendered nearly impossible.

The two tools we developed to enable propagation of cell types from known to unknown organisms are the Nextflow pipeline `kmermaid,` and the Python package `orpheum`. `Kmermaid` addresses the challenge of orthologous gene assignment. `Orpheum` helps `kmermaid` by first detecting putative protein-coding sequence, across ~100 million years of divergence. We benchmarked the prediction of protein-coding sequence using simulated human RNA-seq data from single-copy genes present across all clades of mammals and curated orthologous protein sequences (Simão *et al*., 2015; Altenhoff *et al*., 2016). Given a reference proteome ~100 million years diverged from the organism in question, `orpheum` is able to accurately reject unlikely reading frames. Thus, using a mammalian protein reference database, `orpheum` can unlock the potential of single-cell RNA-seq atlases for any placental mammal.

As confirmation of our methods, we propagated cell types within species (mice) as well as between species (mouse to human, and mouse to bat). Instead of using nucleotide *k*_*nuc*_-mers, our critical innovation is the use of subsampled, translated amino acid *k*_*aa*_-mers. Using *k*_*aa*_-mer signatures, we compressed single cells’ RNA-seq data into subsampled *k*_*aa*_-mers and built cell type databases using sequence bloom trees (SBTs)(Solomon and Kingsford, 2016). Between mice, the performance of cell type label propagation measured by Adjusted Rand Index (ARI) did not differ greatly between the DNA, Protein, and Dayhoff alphabets (Figure 3B-C, green). To test performance across species, we propagated cell type labels from mouse to human (~90 mya diverged) and mouse to *Rhinolophus sinicus*, Chinese greater horseshoe bat (Ren *et al*., 2020) (~95 mya diverged (*TimeTree* :: *The timescale of life*, no date b)). While cell type label propagation across species was low with an ARI ~ 0.4 amongst all molecular alphabets, cellular compartment predictions across species via the Dayhoff alphabet performed well, with ARI ~0.8 as compared to the DNA or Protein alphabets, ARI ~0.6. Our results are consistent with a recent cell type group label classification benchmarking paper showcasing mouse-human visual cortex classification (Abdelaal *et al*., 2019). As all cell types may not be consistently present across species, granular analyses may not be reasonable. Instead, compartment classification metrics such as with `kmermaid`, can still provide valuable output to aid in understanding species’ cell composition.

Overall, several avenues for future optimizations to the *k*-mer homology methods include 1) the use of ensemble scoring of amino acid alphabets (both degenerate and not) to utilize the strengths of the specificity of the 20-letter protein alphabet, and the sensitivity of the degenerate 6-letter Dayhoff amino acid alphabet. 2) For species that diverged > 100 mya it may be useful to consider only *k*_*aa*_-mers from known cell type specific genes or highly conserved transcription factors. 3) Other degenerate protein alphabets, such as a hydrophobic-polar alphabet (Phillips *et al*., 2012) may be useful for evolutionary distances >100 mya. 4) A further enhancement could be to train deep learning models such as scANVI (Xu *et al*., 2021) or MARS (Brbić *et al*., 2020) on the k-mer abundances. One challenge with applying machine learning methods is that there are orders of magnitude more k-mers (~10,000,000 - 1,000,000,000 features) than genes (~5,000-50,000 features), and as a result, the matrices are even sparser than read counts from scRNA-seq data.

`Kmermaid` removes the orthology inference step, opening up the possibility of finding shared and divergent tissue and cell types across a broad range of species and paving the way for evolutionary analyses of cell types across species. To our knowledge, it is the first tool that bypasses the genome alignment steps for cross-species transcriptome comparison. The potential use cases of `kmermaid` in a *de novo* setting for non-model organisms includes finding similar cell types within an organism and finding similar cell types relative to a reference organism, both without the need for a reference genome or reference transcriptome. As the number of scRNA-seq datasets continues to grow, we expect `kmermaid` to be widely used for identifying cell types in non-model organisms. Our results demonstrate the reference-free method using the *k*-mers from single cells is a novel, annotation-agnostic method for comparing cells across species, helping to build the cell type evolutionary tree of life.

For widespread accessibility, portability, and usage, we implemented `kmermaid` as an in-progress portable, containerized Nextflow (Di Tommaso *et al*., 2017) pipeline following software best practices such as testing and continuous integration from the nf-core framework (Ewels *et al*., 2020): https://github.com/nf-core/kmermaid. We welcome any feedback about potential use cases of the *k*-mer homology methods. We provide pre-built cell type databases of from *Tabula Muris Senis* available at https://osf.io/tjxpn/.

## Acknowledgements

The authors deeply thank Angela Pisco and especially Sandra Schmid for their critical reading of the manuscript. The authors also express gratitude to Lia Prins for critical insights into figure design.

## Tables

**Table 2**

*Table 2: Unification of cell type annotations across three species. First three columns, free annotation in Mouse, Human, and Bat. Fourth column, narrow categorization of cell types, e*.*g. with separate T cell subtypes of CD4+, CD8+, Regulatory T, etc. Fifth column, broader categorization of cell types, e*.*g. unifying all T cell types under a single label, “T cell*.*” Third, compartment-level categorization, e*.*g. combining B cell, Plasma, T cell, Proliferating NK/T, Natural Killer cell, and Natural Killer T cells under a single label, “lymphoid*.*” In this work, we focused on the broad and compartment categorizations*.

**Table 3**

*Table 3: Expression of genes unannotated in the bat genome, detected with Dayhoff* k_aa_*-mers unaligned relative to the bat, but contained in reads aligning to known gene loci in the human genome. First column, cell type label; second column, gene category from Supplementary Figure 6B-E; third column, gene symbol from human alignment; fourth column, number of bat cells expressing one or more* k_aa_*-mers matching to the human gene in the third column; fifth column, total number of bat cells in the cell type indicated in the first column; sixth column, percent of cells with a k*_*aa*_*-mer shared with the human gene indicated in the third column*.

## Supplemental Figure Captions

**Supplemental Figure 1.**
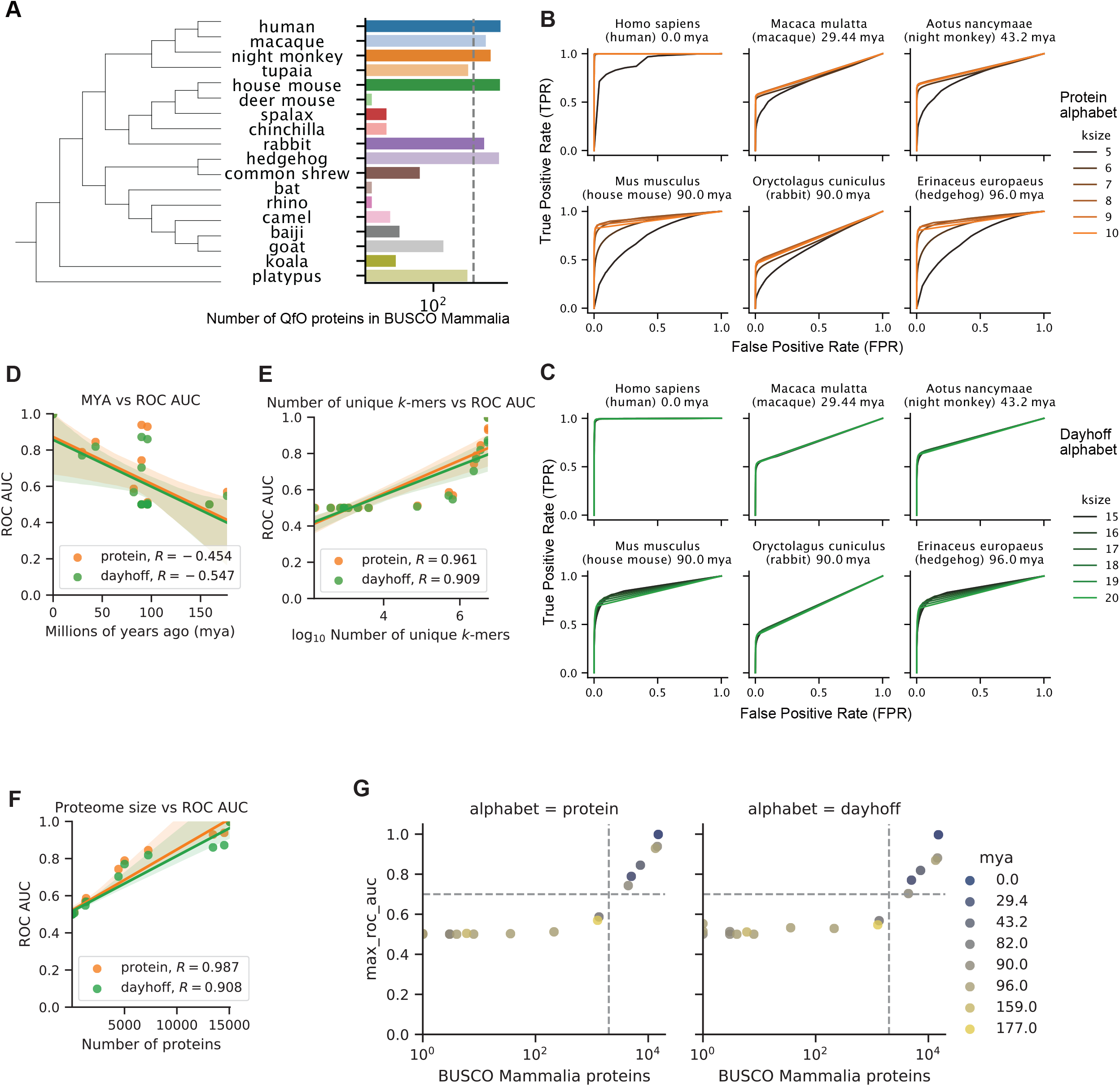
Orpheum detects putative protein-coding sequence across millions of years of evolution. **A**. Phylogenetic tree and proteome sizes of the species in the intersection between the Quest for Orthologs (QfO) 2019 database and BUSCO Mammalia from OrthoDB v10. Left, phylogenetic tree showing evolutionary relationships between mammalian species. Right, barplot of the number of proteins in the organism’s proteome that overlap with BUSCO Mammalia annotations. **B**. ROC curves for performance of `orpheum` using the protein alphabet from proteomes of the six species with minimal 4000 proteins. Only ksize = 5 … 10 is shown as performance declines outside of this range. **C**. ROC curves for performance of `orpheum` using the Dayhoff alphabet from proteomes of the six species with minimal 4000 proteins. Only ksize = 15 … 20 is shown as performance declines outside of this range. **D**. Proteome size is not strongly dependent with divergence time from human. x-axis, millions of years the species is diverged from human, as identified from predicted divergence time from timetree.org. y-axis, number of proteins in the species’ proteome from the Quest for Orthologs (QfO) that match benchmarking universal single copy orthologs (BUSCO) Mammalia annotations. **E**. Predictive power of `orpheum` is strongly dependent on the number of unique *k*-mers in the database. Predictive power of `orpheum` to detect protein-coding sequence across ~200 million years of evolution, using simulated human RNA-seq reads and mammalian reference proteomes. x-axis, *k*-mer size used to translate the RNA sequence; y-axis, receiver operating characteristic area under the curve (ROC AUC) of protein-coding prediction at that *k*-mer size. Dotted vertical lines indicate *k*-mer size producing maximum ROC AUC for each divergence time. **F**. Predictive power of `orpheum` is strongly dependent on the number of proteins present in the protein database. Millions of years of divergence compared to ROC AUC. Spearman’s rank-based correlation coefficient is shown. **G**. Predictive power of `orpheum` depends on the input proteome size, not on the divergence time from the query species. x-axis, number of species’ proteins present in BUSCO Mammalia annotation, y-axis, maximum value of ROC AUC across all *k*_*aa*_-mer sizes and amino acid alphabets. Dotted line: 4000 proteins on x-axis and 0.7 ROC AUC value.

**Supplemental Figure 2.**
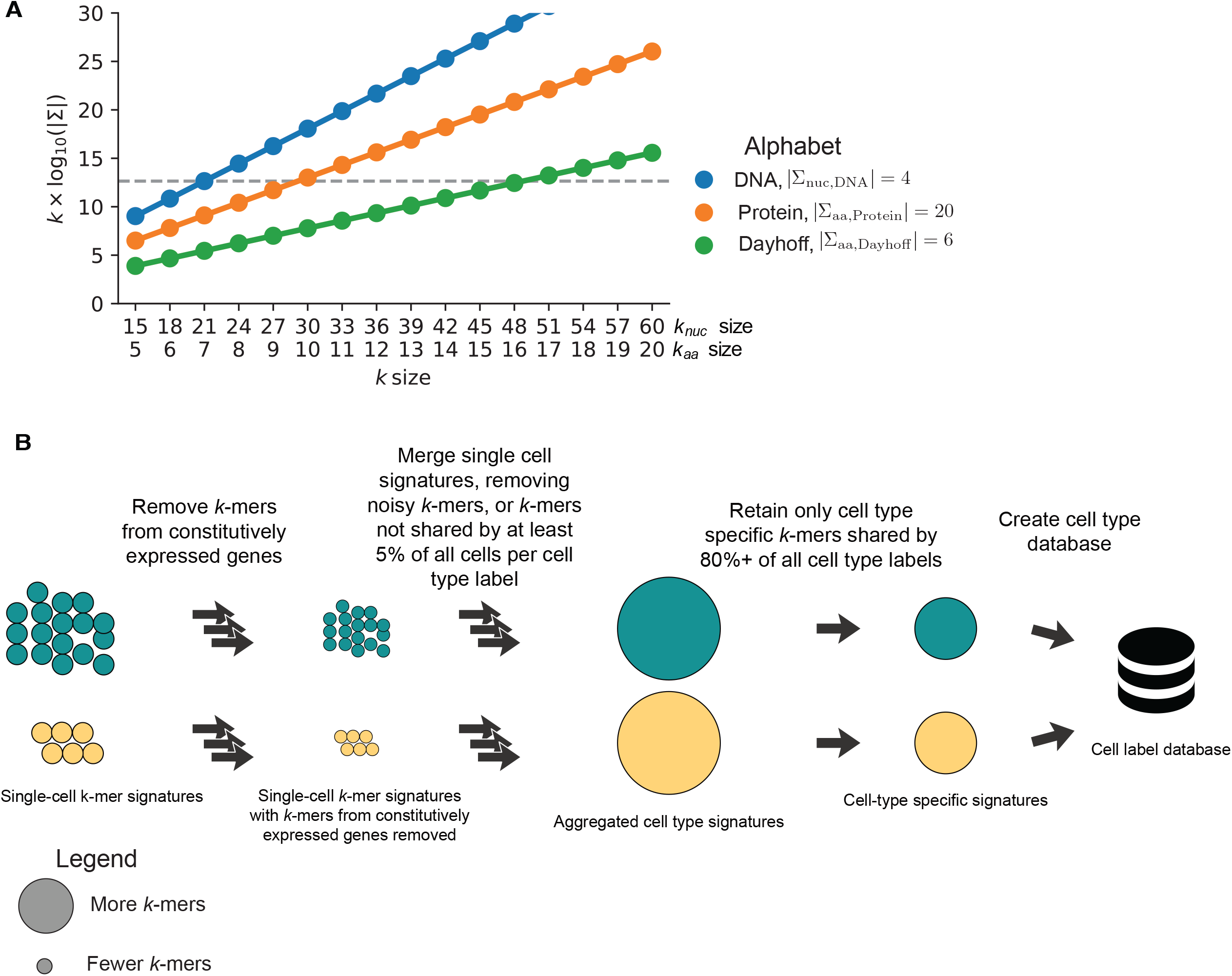
Amino acid *k*_*aa*_-size determination and cell type database construction. **A**. Comparison of complexity of nucleotide (4-letter), protein (20-letter), and degenerate amino acid Dayhoff (6-letter) alphabets. X-axis, *k*_*nuc*_- and *k*_*aa*_-mer sizes, and y-axis, log_10_-scaled number of *k*-mers present at that k-mer size for each alphabet. Blue, DNA, orange, protein, green, Dayhoff alphabets. **B**. Cell type database construction from single cells.

**Supplemental Figure 3.**
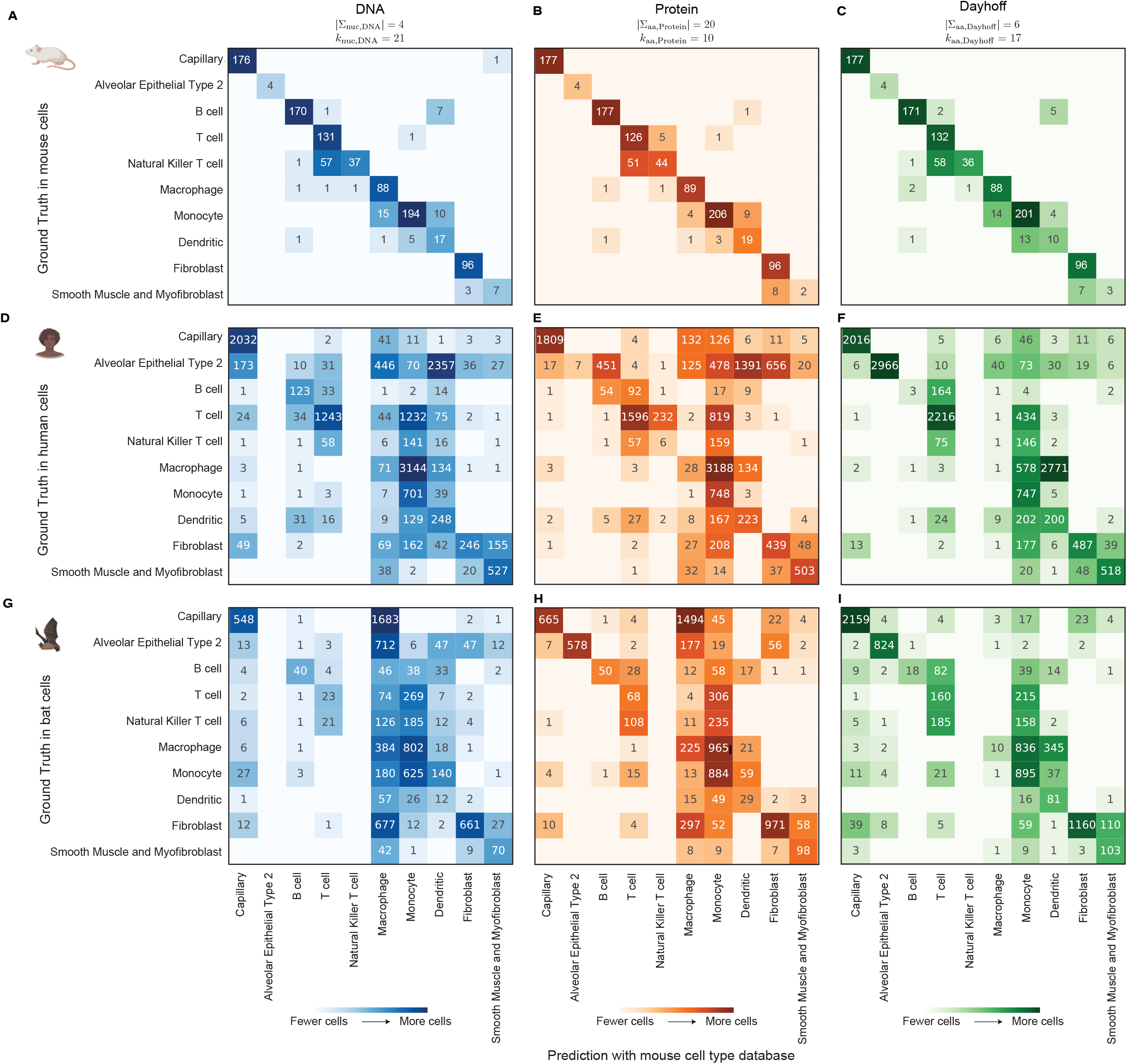
Confusion matrices of cell type-level classification from a mouse database. Rows, ground truth label and columns, predicted label. Numbers in the matrices indicate the number of cells in the ground truth label, to be labeled as the predicted label. **A-C**: Mouse to mouse mapping, using a cell type database from one mouse, and classifying cell types with a different mouse. **A**. Classification using DNA *k*_*nuc*_-mers (|∑_nuc, DNA_| = 4, *k*_*DNA*_ = 21). **B**. Classification using protein *k*_*aa*_-mers (|∑_aa, Protein_, DNA| = 20, *k*_*Protein*_ = 10). **C**. Classification using Dayhoff *k*_*aa*_-mers (|∑_aa, Dayhoff_| = 6, *k*_*Dayhoff*_ = 17). **D-F**: Mouse to human mapping, using a cell type database from many mice, and classifying cell types in human single cells. **D**. Classification using DNA *k*_*nuc*_-mers (|∑_nuc, DNA_| = 4, *k*_*DNA*_ = 21). **E**. Classification using protein *k*_*aa*_-mers (|∑_aa, Protein_, DNA| = 20, *k*_*Protein*_ = 10). **F**. Classification using Dayhoff *k*_*aa*_-mers (|∑_aa, Dayhoff_| = 6, *k*_*Dayhoff*_ = 17). **G-I**: Mouse to bat mapping, using a cell type database from many mice, and classifying cell types in bat single cells. **G**. Classification using DNA *k*_*nuc*_-mers (|∑_nuc, DNA_| = 4, *k*_*DNA*_ = 21). **H**. Classification using protein *k*_*aa*_-mers (|∑_aa, Protein_, DNA| = 20, *k*_*Protein*_ = 10). **I**. Classification using Dayhoff *k*_*aa*_-mers (|∑_aa, Dayhoff_| = 6, *k*_*Dayhoff*_ = 17).

**Supplemental Figure 4.**
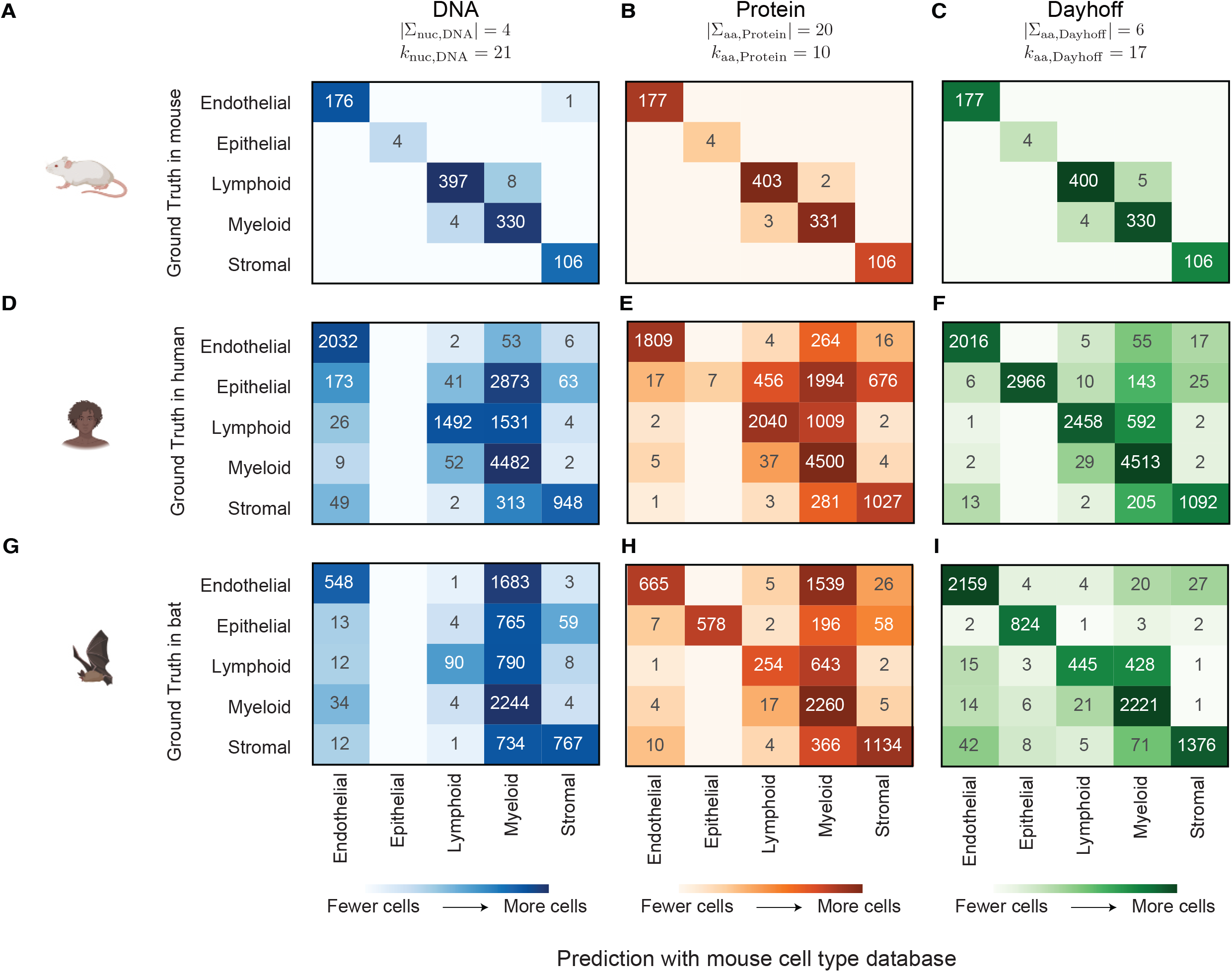
Confusion matrices of cellular compartment-level classification from a mouse database. Rows, ground truth label and columns, predicted label. Rows, ground truth label and columns, predicted label. Numbers in the matrices indicate the number of cells in the ground truth label, to be labeled as the predicted label. **A-C**: Mouse to mouse mapping, using a cell type database from one mouse, classifying cell types with a different mouse, and aggregating to cellular compartments. **A**. Classification using DNA *k*_*nuc*_-mers (|∑_nuc, DNA_| = 4, *k*_*DNA*_ = 21). **B**. Classification using protein *k*_*aa*_-mers (|∑_aa, Protein_, DNA| = 20, *k*_*Protein*_ = 10). **C**. Classification using Dayhoff *k*_*aa*_-mers (|∑_aa, Dayhoff_| = 6, *k*_*Dayhoff*_ = 17). **D-F**: Mouse to human mapping, using a cell type database from many mice, and classifying cell types in human single cells, and aggregating to cellular compartments. **D**. Classification using DNA *k*_*nuc*_-mers (|∑_nuc, DNA_| = 4, *k*_*DNA*_ = 21). **E**. Classification using protein *k*_*aa*_-mers (|∑_aa, Protein_, DNA| = 20, *k*_*Protein*_ = 10). **F**. Classification using Dayhoff *k*_*aa*_-mers (|∑_aa, Dayhoff_| = 6, *k*_*Dayhoff*_ = 17). **G-I**: Mouse to bat mapping, using a cell type database from many mice, and classifying cell types in bat single cells, and aggregating to cellular compartments. **G**. Classification using DNA *k*_*nuc*_-mers (|∑_nuc, DNA_| = 4, *k*_*DNA*_ = 21). **H**. Classification using protein *k*_*aa*_-mers (|∑_aa, Protein_, DNA| = 20, *k*_*Protein*_ = 10). **I**. Classification using Dayhoff *k*_*aa*_-mers (|∑_aa, Dayhoff_| = 6, *k*_*Dayhoff*_ = 17).

**Supplemental Figure 5.**
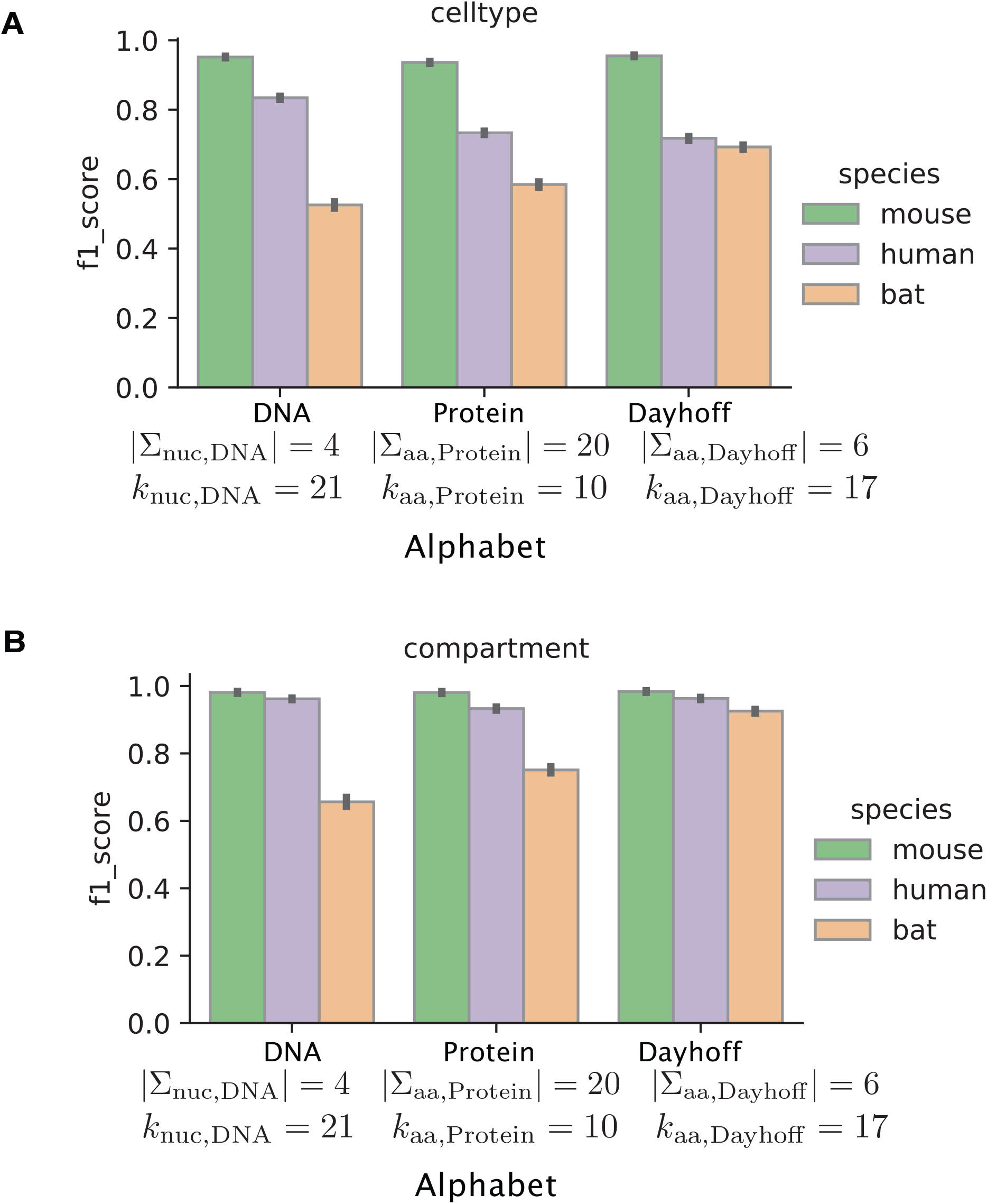
**A**. F1 scores for cell type classification across DNA, protein, and Dayhoff alphabets, for the three species of mouse, human and bat. **B**. F1 scores for cellular compartment classification across DNA, protein, and Dayhoff alphabets, for the three species of mouse, human and bat.

**Supplemental Figure 6.**
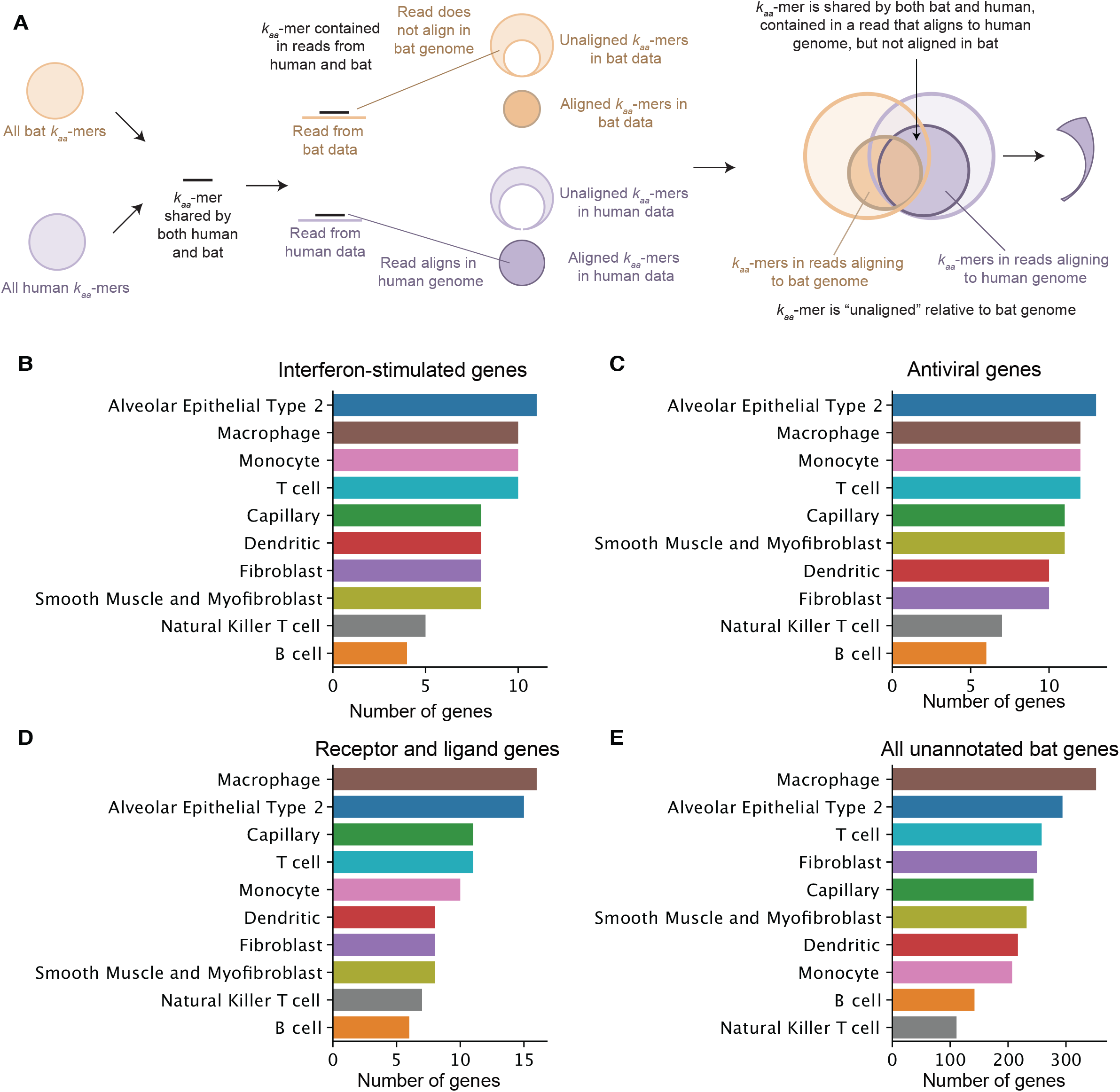
Unaligned Dayhoff *k*_*aa*_-mers in bat are contained in human reads aligning to known cell type marker genes. **A**. Schematic demonstrating an “unaligned *k*-mer” in the bat genome, but this *k*-mer was contained in a read that aligned to the human genome. **B**. Number of interferon-stimulated genes (ISGs) identified per ground truth cell type from unaligned Dayhoff *k*_*aa*_-mers in bat, matching to known genes in human. **C**. Number of antiviral genes identified per ground truth cell type from unaligned Dayhoff *k*_*aa*_-mers in bat, matching to known genes in human. **D**. Number of receptor-ligand genes identified per ground truth cell type from unaligned Dayhoff *k*_*aa*_-mers in bat, matching to known genes in human. **E**. Number of human genes unannotated in the *R. sinicus* genome, identified per ground truth cell type from unaligned Dayhoff *k*_*aa*_-mers in bat.

## Notes

https://github.com/nf-core/kmermaid

https://github.com/czbiohub/orpheum

https://osf.io/tjxpn/

### Competing Interest Statement

The authors have declared no competing interest.

## References

Abdelaal, T. et al. (2019) ‘A comparison of automatic cell identification methods for single-cell RNA sequencing data’, Genome biology, 20(1), p. 194. doi: 10.1186/s13059-019-1795-z.

Altenhoff, A. M. et al. (2016) ‘Standardized benchmarking in the quest for orthologs’, Nature methods, 13(5), pp. 425–430. doi: 10.1038/nmeth.3830.

Altenhoff, A. M. et al. (2020) ‘The Quest for Orthologs benchmark service and consensus calls in 2020’, Nucleic acids research, 48(W1), pp. W538–W545. doi: 10.1093/nar/gkaa308.

Brbić, M. et al. (2020) ‘MARS: discovering novel cell types across heterogeneous single-cell experiments’, Nature methods. doi: 10.1038/s41592-020-00979-3.

Brown, C. T. and Irber, L. (2016) ‘sourmash: a library for MinHash sketching of DNA’, The Journal of Open Source Software, 1(5), p. 27. doi: 10.21105/joss.00027.

Buchfink, B., Xie, C. and Huson, D. H. (2015) ‘Fast and sensitive protein alignment using DIAMOND’, Nature methods, 12(1), pp. 59–60. doi: 10.1038/nmeth.3176.

Compeau, P. and Pevzner, P. A. (2018) Bioinformatics Algorithms: An Active Learning Approach. La Jolla. CA: Active Learning Publishers. Available at: https://fokt.pw/zora-nog-ry-k.pdf.

DAYHOFF and M. O (1972) ‘A model of evolutionary change in proteins’, Atlas of Protein Sequence and Structure, 5, pp. 89–99. Available at: https://ci.nii.ac.jp/naid/10024396603/ (Accessed: 19 December 2020).

Dayhoff, M. O. (1972) Atlas of protein sequence and structure. National Biomedical Research Foundation.

Dayhoff, M. O. and National Biomedical Research Foundation (1969) Atlas of protein sequence and structure: 1969. National Biomedical Research Foundation. Available at: https://play.google.com/store/books/details?id=9ODSxQEACAAJ.

Di Tommaso, P. et al. (2017) ‘Nextflow enables reproducible computational workflows’, Nature biotechnology, 35(4), pp. 316–319. doi: 10.1038/nbt.3820.

Edgar, R. C. (2004) ‘Local homology recognition and distance measures in linear time using compressed amino acid alphabets’, Nucleic acids research, 32(1), pp. 380–385. doi: 10.1093/nar/gkh180.

Emms, D. M. and Kelly, S. (2015) ‘OrthoFinder: solving fundamental biases in whole genome comparisons dramatically improves orthogroup inference accuracy’, Genome biology, 16, p. 157. doi: 10.1186/s13059-015-0721-2.

Emms, D. M. and Kelly, S. (2019) ‘OrthoFinder: phylogenetic orthology inference for comparative genomics’, Genome biology, 20(1), p. 238. doi: 10.1186/s13059-019-1832-y.

Ewels, P. A. et al. (2020) ‘The nf-core framework for community-curated bioinformatics pipelines’, Nature biotechnology, 38(3), pp. 276–278. doi: 10.1038/s41587-020-0439-x.

da Fonseca, R. R. et al. (2016) ‘Next-generation biology: Sequencing and data analysis approaches for non-model organisms’, Marine genomics, 30, pp. 3–13. doi: 10.1016/j.margen.2016.04.012.

Formenti, G. et al. (2020) ‘Complete vertebrate mitogenomes reveal widespread gene duplications and repeats’, Cold Spring Harbor Laboratory. doi: 10.1101/2020.06.30.177956.

Hampikian, G. and Andersen, T. (2006) ‘ABSENT SEQUENCES: NULLOMERS AND PRIMES’, in Biocomputing 2007. WORLD SCIENTIFIC, pp. 355–366. doi: 10.1142/9789812772435_0034.

Hastie, T., Tibshirani, R. and Friedman, J. (2009) The Elements of Statistical Learning: Data Mining, Inference, and Prediction, Second Edition (Springer Series in Statistics). Available at: https://www.amazon.com/Elements-Statistical-Learning-Prediction-Statistics/dp/0387848576.

Huerta-Cepas, J. et al. (2016) ‘eggNOG 4.5: a hierarchical orthology framework with improved functional annotations for eukaryotic, prokaryotic and viral sequences’, Nucleic acids research, 44(D1), pp. D286–D293. Available at: https://academic.oup.com/nar/article-lookup/doi/10.1093/nar/gkv1248.

Huerta-Cepas, J. et al. (2017) ‘Fast Genome-Wide Functional Annotation through Orthology Assignment by eggNOG-Mapper’, Molecular biology and evolution, 34(8), pp. 2115–2122. Available at: https://academic.oup.com/mbe/article/34/8/2115/3782716.

Hu, X. and Friedberg, I. (2019) ‘SwiftOrtho: A fast, memory-efficient, multiple genome orthology classifier’, GigaScience, 8(10), pp. 309–312. Available at: https://academic.oup.com/gigascience/article/doi/10.1093/gigascience/giz118/5606727.

Irber, L. and Brown, C. T. (2020) ‘Lightweight compositional analysis of metagenomes with sourmash gather’, Manubot. Available at: https://dib-lab.github.io/2020-paper-sourmash-gather/ (Accessed: 16 December 2020).

Landès, C. and Risler, J. L. (1994) ‘Fast databank searching with a reduced amino-acid alphabet’, Computer applications in the biosciences: CABIOS, 10(4), pp. 453–454. doi: 10.1093/bioinformatics/10.4.453.

Liu, X. et al. (2011) ‘Requirement of the nuclear localization of transcription enhancer factor 3 for proliferation, migration, tube formation, and angiogenesis induced by vascular endothelial growth factor’, FASEB journal: official publication of the Federation of American Societies for Experimental Biology, 25(4), pp. 1188–1197. doi: 10.1096/fj.10-167619.

MGI-Mouse Genome Informatics-The international database resource for the laboratory mouse (no date). Available at: http://www.informatics.jax.org/ (Accessed: 1 July 2021).

Miga, K. H. et al. (2020) ‘Telomere-to-telomere assembly of a complete human X chromosome’, Nature, 585(7823), pp. 79–84. doi: 10.1038/s41586-020-2547-7.

Mora, C. et al. (2011) ‘How Many Species Are There on Earth and in the Ocean?’, PLoS biology, 9(8), pp. e1001127–8. Available at: https://dx.plos.org/10.1371/journal.pbio.1001127.

Murphy, L. R., Wallqvist, A. and Levy, R. M. (2000) ‘Simplified amino acid alphabets for protein fold recognition and implications for folding’, Protein engineering, 13(3), pp. 149–152. doi: 10.1093/protein/13.3.149.

Neff, N. F. et al. (2018) ‘Single-cell transcriptomics of 20 mouse organs creates a Tabula Muris’, Nature, pp. 1–25. doi: 10.1038/s41586-018-0590-4.

Nichio, B. T. L., Marchaukoski, J. N. and Raittz, R. T. (2017) ‘New Tools in Orthology Analysis: A Brief Review of Promising Perspectives’, Frontiers in genetics, 8, p. 165. doi: 10.3389/fgene.2017.00165.

Ondov, B. D. et al. (2016) ‘Mash: fast genome and metagenome distance estimation using MinHash’, Genome biology, pp. 1–14. doi: 10.1186/s13059-016-0997-x.

Ondov, B. D. et al. (2019) ‘Mash Screen: high-throughput sequence containment estimation for genome discovery’, Genome biology, 20(1), pp. 1–13. doi: 10.1186/s13059-019-1841-x.

Peris, P., López, D. and Campos, M. (2008) ‘IgTM: An algorithm to predict transmembrane domains and topology in proteins’, BMC bioinformatics, 9(1), pp. 1029–1011. Available at: https://bmcbioinformatics.biomedcentral.com/articles/10.1186/1471-2105-9-367.

Peterson, E. L. et al. (2009) ‘Reduced amino acid alphabets exhibit an improved sensitivity and selectivity in fold assignment’, Bioinformatics, 25(11), pp. 1356–1362. doi: 10.1093/bioinformatics/btp164.

Phillips, R. et al. (2012) Physical Biology of the Cell. Garland Science. Available at: http://books.google.com/books?id=5_MOBAAAQBAJ&printsec=frontcover&dq=intitle:Physical+Biology+of+the+Cell&hl=&cd=1&source=gbs_api.

Pierce, N. T. et al. (2019) ‘Large-scale sequence comparisons with sourmash’, F1000Research, 8, p. 1006. doi: 10.12688/f1000research.19675.1.

Regev, A. et al. (2017) ‘The Human Cell Atlas’, pp. 1–61. Available at: http://biorxiv.org/lookup/doi/10.1101/121202.

Ren, L. et al. (2020) ‘Single-cell transcriptional atlas of the Chinese horseshoe bat (Rhinolophus sinicus) provides insight into the cellular mechanisms which enable bats to be viral reservoirs’, 6, pp. 23–62. Available at: http://biorxiv.org/lookup/doi/10.1101/2020.06.30.175778.

Rhie, A. et al. (2020) ‘Towards complete and error-free genome assemblies of all vertebrate species’, Cold Spring Harbor Laboratory. doi: 10.1101/2020.05.22.110833.

Scheffers, B. R. et al. (2012) ‘What we know and don’t know about Earth’s missing biodiversity’, Trends in ecology & evolution, 27(9), pp. 501–510. doi: 10.1016/j.tree.2012.05.008.

Shafer, M. E. R. (2019) ‘Cross-Species Analysis of Single-Cell Transcriptomic Data’, Frontiers in cell and developmental biology, 7, p. 175. doi: 10.3389/fcell.2019.00175.

Shi, C. H. and Yip, K. Y. (2019) ‘K-mer counting with low memory consumption enables fast clustering of single-cell sequencing data without read alignment’, Cold Spring Harbor Laboratory. doi: 10.1101/723833.

Simão, F. A. et al. (2015) ‘BUSCO: assessing genome assembly and annotation completeness with single-copy orthologs’, Bioinformatics, 31(19), pp. 3210–3212. doi: 10.1093/bioinformatics/btv351.

Solomon, B. and Kingsford, C. (2016) ‘Fast search of thousands of short-read sequencing experiments’, Nature biotechnology, 34(3), pp. 300–302. doi: 10.1038/nbt.3442.

Sonnhammer, E. L. L. et al. (2014) ‘Big data and other challenges in the quest for orthologs’, Bioinformatics, 30(21), pp. 2993–2998. doi: 10.1093/bioinformatics/btu492.

Tabula Muris Consortium (2020) ‘A single-cell transcriptomic atlas characterizes ageing tissues in the mouse’, Nature, 583(7817), pp. 590–595. doi: 10.1038/s41586-020-2496-1.

Tarashansky, A. J. et al. (2021) ‘Mapping single-cell atlases throughout Metazoa unravels cell type evolution’. doi: 10.7554/eLife.66747.

TimeTree :: The timescale of life (no date a). Available at: http://timetree.org (Accessed: 21 December 2020).

TimeTree :: The timescale of life (no date b). Available at: http://timetree.org (Accessed: 24 February 2021).

Travaglini, K. J. et al. (2019) ‘A molecular cell atlas of the human lung from single cell RNA sequencing’, bioRxiv, 79(1), p. 742320. Available at: http://biorxiv.org/lookup/doi/10.1101/742320.

Vilella, A. J. et al. (2008) ‘EnsemblCompara GeneTrees: Complete, duplication-aware phylogenetic trees in vertebrates’, Genome research, 19(2), pp. 327–335. Available at: http://genome.cshlp.org/cgi/doi/10.1101/gr.073585.107.

Xu, C. et al. (2021) ‘Probabilistic harmonization and annotation of single-cell transcriptomics data with deep generative models’, Molecular systems biology, 17(1), p. e9620. doi: 10.15252/msb.20209620.

Yates, A. D. et al. (2020) ‘Ensembl 2020’, Nucleic acids research, 48(D1), pp. D682–D688. doi: 10.1093/nar/gkz966.

Ye, Y., Choi, J.-H. and Tang, H. (2011) ‘RAPSearch: a fast protein similarity search tool for short reads’, BMC bioinformatics, 12, p. 159. doi: 10.1186/1471-2105-12-159.

