## Supplementary Methods for "Single-cell transcriptomics for the 99.9% of species without reference genomes"

*Olga Botvinnik*

*July 9, 2021*

### *Contents*

*Orpheum algorithmic details* 2

*Sequence complexity filters* 2

*Reduced alphabets* 2

*False Positive Rate in* `orpheum translate` 2

*Benchmarking dataset creation* 6

*Installation* 6

*Read preprocessing before* `orpheum translate` 6

*Usage* 6

`kmermaid` *pipeline* 8

*Comment on* `kmermaid` *usage on organisms without reference genomes* 9

*Sequence data processing* 10

*Datasets* 10

*Cell type database creation* 10

*Removal of k-mers uninformative for cell type annotations* 10

*Merging of single cell signatures and removal of "noisy" k-mers* 11

*Removal of non cell type-specific k-mers* 11

*Mouse to mouse database creation and cell type testing* 11

*Mouse cell type database creation for cross-species* 11

### *Orpheum algorithmic details*

Orpheum finds putative protein-coding sequences by performing six-frame translation of RNA-seq reads.

#### *Sequence complexity filters*

To provide high-quality protein-coding predictions, we filter each reading frame's translated amino acid seuqence for sequence complexity in two ways:

1. Consecutive amino acid sequence complexity, using the same formula as in the fastp tool[1] for sequence processing, and a threshold of 0.3.

2. Given a reading frame of length $L_{\text{translated}}$, the number of unique $k$-mers per reading frame must be greater than or equal to $\frac{L_{\text{translated}}-k+1}{2}$

[1] Chen et al. 2018.

#### *Reduced alphabets*

At the core of `orpheum` is the ability to cheaply compare sequences using short, $k$-long substrings, or $k$-mers. However, a single residue change in a $k$-mer changes the whole sequence, and thus $k$-mers are brittle to single-residue substitutions. To compare across evolutionary distances, where small amino acid changes may occur that do not drastically change the resulting protein, one must allow for minor base substitutions that still maintain similar chemical or structural properties. A reduced alphabet can encode useful information into a smaller alphabet space, and enable sequence comparisons across a broader variety of species than the original alphabet alone.

```
protein20: SASHAFIERCE
dayhoff6:  bbbdbfecdac
```

Figure 1: Example of the Dayhoff reduced amino alphabet applied to the amino acid sequence, `SASHAFIERCE`.

*Reduced amino acid alphabets* Reduced amino acid alphabets have a greater than 50-year history[2] in finding related protein sequences.[3] Recently, a reduced amino acid alphabet combined with $k$-mers were used to find homologous protein-coding sequences.[4] We build on this concept by enabling prediction of protein-coding sequences from RNA-seq reads using Protein and Dayhoff alphabets, shown by Table 1 in main text and Figure 1.

[2] Dayhoff 1969.

[3] Edgar 2004; Murphy et al. 2000; Peterson et al. 2009; Landès et al. 1994; Ye et al. 2011.

[4] Hu et al. 2019.

#### *False Positive Rate in `orpheum translate`*

`orpheum translate` finds putative protein-coding frames performing six-frame translation and computing the $k$-mer containment between each reading frame and the proteome database.

Ultimately, we chose a containment threshold of at least 5% of $k$-mers from the reading frame must overlap with the query sequence.

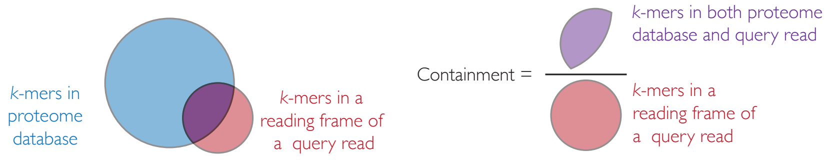


Figure 2: Illustration of $k$-mer containment computed between putative protein-coding reading frames and the proteome database.

To set a threshold of the minimum Jaccard overlap between a translated read's frame and the reference proteome, the most statistically principled way is to control the false positive rate (FPR) of predicting a protein-coding read, which we compute below.

*Probability of random k-mers from a read* If $k$-mers from reads were independent, identically distributed (i.i.d.) variables, then a translated read of length $L_{\text{translated}}$ drawing letters from the alphabet $\Sigma$, whose size is $|\Sigma|$, would contain at most

$$\left(\frac{1}{|\Sigma|^k}\right)^{L_{\text{translated}}-k+1} \tag{1}$$

$k$-mers per read.

However, $k$-mers drawn from reads are not i.i.d. Let's take a simple example. If we have a two-letter alphabet, $\Sigma = \{a, b\}$, thus $|\Sigma| = 2$. Let us use an example sequence $S = abbabba$. If $k = 4$, then the first $k$-mer is $abba$. The second $k$-mer is thus either $bbaa$ or $bbab$, with equal probability. We can generalize this: Given the first $k$-mer, the first $k-1$ letters from the second $k$-mer are known, and thus the probability of guessing the next $k$-mer is $\frac{1}{|\Sigma|}$.

Thus, the probability of a random $k$-mer from a sequencing read is completely dependent on the alphabet size $|\Sigma|$ and its translated sequence length $L_{\text{translated}}$ is

$$\Pr(\text{FPR}_{k-\text{mer}}) = \left(\frac{1}{|\Sigma|}\right)^{k} \times \left(\frac{1}{|\Sigma|}\right)^{L_{\text{translated}}-k} \tag{2}$$

$$= \left(\frac{1}{|\Sigma|}\right)^{L_{\text{translated}}-k+k} \tag{3}$$

$$= \left(\frac{1}{|\Sigma|}\right)^{L_{\text{translated}}}. \tag{4}$$

$$= |\Sigma|^{-L_{\text{translated}}} \tag{5}$$

Note that this is equal to the inverse of the number of sequences of length $L_{\text{translated}}$ with alphabet size $|\Sigma|$, or $|\Sigma|^{L_{\text{translated}}}$.

*Bloom filter collision probability* A Bloom Filter[5] is a probabalistic

[5] Leskovec et al. 2020.

data structure that efficiently stores set membership. The tradeoffs of a Bloom filter include very efficient memory and time space to test membership with no false negatives, at the cost of some false positives. The probability of error of the Bloom filter implementation in `khmer`[6] used in `orpheum`, given $N$ distinct $k$-mers counted, a hash table size of $H$, and $Z$ total number of hash tables, is

[6] Crusoe et al. 2015.

$$\Pr(\text{FPR}_{\text{bloom}}) = \left(1 - \left(1 - \frac{1}{H \times Z}\right)^{ZN}\right)^{Z} \tag{6}$$

$$\approx \left(1 - \exp\left(-\frac{NZ}{HZ}\right)\right)^{Z} \tag{7}$$

$$= \left(1 - \exp\left(-\frac{N}{H}\right)\right)^{Z}, \tag{8}$$

where the total size of the Bloom filter, often called $m$, is approximately[7] $m \approx H \times Z$.

[7] See the discussion on the thread here: dib-lab/khmer#1898

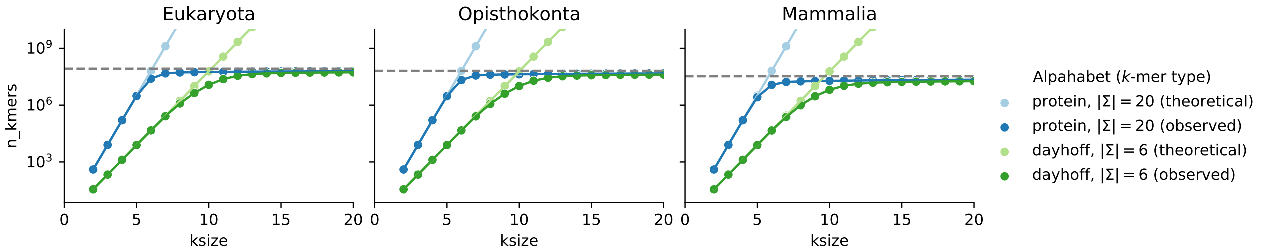


Figure 3: Number of amino acid $k$-mers observed in UniProt/SwissProt manually curated databases. Each dataset was manually downloaded from uniprot.org, then unique $k$-mers at each amino acid $k$-mer size was computed using the Bloom Filter implementation in `khmer.NodeGraph`.

Theoretically, the total number of $k$-mers is limited by the alphabet size and choice of $k$. Empirically, the number of possible $k$-mers is limited by the $k$-mers observed in the databases. As shown in Figure 3, by $k = 5$, the number of theoretical protein $k$-mers exceeds the number of observed protein $k$-mers. Additionally, the mass of all possible DNA $k$-mers of length 79 or more exceeds the mass of the earth,[8] and thus it is impossible to observe all theoretical amino acid $k$-mers beyond length $\frac{79}{3} = 26$. The largest database tested, the UniProtKB Eukaryota manually reviewed dataset contains $5 \times 10^7$ 7-mers in the protein alphabet. Thus, we can give an upper bound to the number of possible $k$-mers to be $5 \times 10^7$. Therefore, we approximate the total number of $k$-mers in the bloom filter as,

[8] Louis 2016.

$$N = \min\left(5 \times 10^7, |\Sigma|^k\right) \tag{9}$$

$$= 5 \times 10^7, \forall k_{aa,\text{Protein}} > 5, \tag{10}$$

where $k_{aa,\text{Protein}}$ indicates the $k$-mer size for amino acids in the Protein alphabet.

*False positive rate of protein-coding prediction* Combining Equations (5), (1), (10), and (8) with RNA-seq read of length $L$ where its translated length $L_{\text{translated}} = \lfloor \frac{L}{3} \rfloor$, containing a possible six frames of translation, then the false-positive rate (FPR) protein-coding read is,

$$\Pr\left(\text{FPR}_{k\text{-mer in bloom filter}}\right) = \Pr(\text{FPR}_{k-\text{mer}}) \times \Pr(\text{FPR}_{\text{bloom}}) \tag{11}$$

$$= 6 \times \left(1 - \exp\left(-\frac{N}{H}\right)\right)^{Z} \times |\Sigma|^{-L_{\text{translated}}}. \tag{12}$$

In our implementation, we let the number of possible $k$-mers be $5 \times 10^7$, the number of hash tables $Z = 4$, each table size is $H = 10^8$. Let us assume a read length $L = 90$ (a lower bound given modern short read sequencing technologies with read lengths of 150), where the longest translated frame is $L_{\text{translated}} = 30$. Finally, let us use the Protein alphabet, where $|\Sigma| = 20$. Then, the equation becomes

$$\Pr\left(\text{FPR}_{k\text{-mer in bloom filter}}\right) = 6 \times \left(1 - \exp\left(-\frac{5 \times 10^7}{10^8}\right)\right)^4 \times 20^{-30}. \tag{13}$$

$$\log \Pr\left(\text{FPR}_{k\text{-mer in bloom filter}}\right) = \log(6) + 4\log\left(1 - \exp\left(-\frac{1}{2}\right)\right) - 30\log(20) \tag{14}$$

$$\approx -91.8, \tag{15}$$

an extremely low probability of false positive protein coding sequences.

However, Equation (15) was calculated if ALL the $k$-mers from the protein coding frame are present. In reality, we use a threshold of at least 5% of $k$-mers present in the proteome. This changes the equation to:

$$\log \Pr\left(\text{FPR}_{k\text{-mer in bloom filter}}\right) = \log(6) + 4\log\left(1 - \exp\left(-\frac{1}{2}\right)\right) - 0.5 \times 30\log(20) \tag{16}$$

$$\approx -6.43 \tag{17}$$

Thus,

$$\Pr\left(\text{FPR}_{k\text{-mer in bloom filter}}\right) = \exp\left(-6.43\right) \tag{18}$$

$$= 0.00161, \tag{19}$$

or approximately 1/1000 reading frames may be incorrectly called as protein-coding. As we saw in the main text's Figure 2E-F, the precision of rejecting incorrect reading frames was nearly perfect, and thus we are confident that even this small false positive rate will not strongly affect our results.

### *Benchmarking dataset creation*

For benchmarking, we unified the identifiers of genes from BUSCO Mammalia[9] and the Quest for Orthologs.[10] To create a strict dataset of known orthologs across species, we used a threshold of at least 4000 proteins present in both BUSCO Mammalia and the Quest for Orthologs 2019 reference proteome datasets.

[9] Waterhouse et al. 2018; Simão et al. 2015.

[10] Sonnhammer et al. 2014; Altenhoff et al. 2020; Capella-Gutierrez et al. 2017; Glover et al. 2019.

### *Installation*

`orpheum` can be installed with the Python Package Index (PyPI) with the Python package manager, `pip`, e.g. with the command

```
pip install orpheum
```

### *Read preprocessing before `orpheum translate`*

As the protein-coding score is assessed on the entire read, we recommend RNA-seq reads be removed of library artifacts to the best of the user's ability. This means, the adapters should be trimmed, and if there was a negative insert size such that the R1 and R2 reads overlap, then the read pairs should be merged.

### *Usage*

*Creation of amino acid k-mer database with `orpheum index`* Before predicting protein-coding sequences, `orpheum` must create a database of known amino acid *k*-mers, which is stored in the form of a probabilistic set membership data structure known as a bloom filter. `orpheum` uses the bloom filter implementation in `khmer/oxli`,[11] called a `NodeGraph`. We created a dataset of known amino acid *k*-mers from the manually annotated UniProtKB/Swiss-Prot databases.[12] We used only protein sequences observed in *Mammalia* species.

[11] Crusoe et al. 2015; Zhang et al. 2014.

[12] Consortium 2020; Boutet et al. 2016.

The following command:

```
orpheum index mammalia.fasta.gz
```

will create a proteome database with the Protein alphabet and $k_{aa} = 8$, saving as the file:
`mammalia.fasta.gz.alphabet-protein_ksize-8.bloomfilter.nodegraph`.

*Prediction of protein-coding sequences with `orpheum translate`* We then predicted protein coding reads using the created bloom filter using `orpheum translate`, with the following command:

```
orpheum translate \
  --molecule protein \
  --coding-nucleotide-fasta sample__coding_reads_nucleotides.fasta \
  --csv sample__coding_scores.csv \
```

```
--json-summary sample__coding_summary.json \
--peptides-are-index \
mammalia.fasta.gz.alphabet-protein\_ksize-8.bloomfilter.nodegraph \
sample_R1.fastq.gz sample_R2.fastq.gz \
> sample__coding_reads_peptides.fasta
```

Note that the protein-coding reads are sent to standard output by default, which can help with using Unix pipes between programs.

### `kmermaid` *pipeline*

The `kmermaid` pipeline creates *k*-mer signatures from sequence data. If the provided data is a `.bam` alignment file or a `.tgz` file containing the output from 10x Genomics' `cellranger` program, then we use bam2fasta[13] and Samtools[14] to extract per-cell fastq sequence files.

[13] *bam2fasta* n.d.

[14] Danecek et al. 2021; Li et al. 2009.

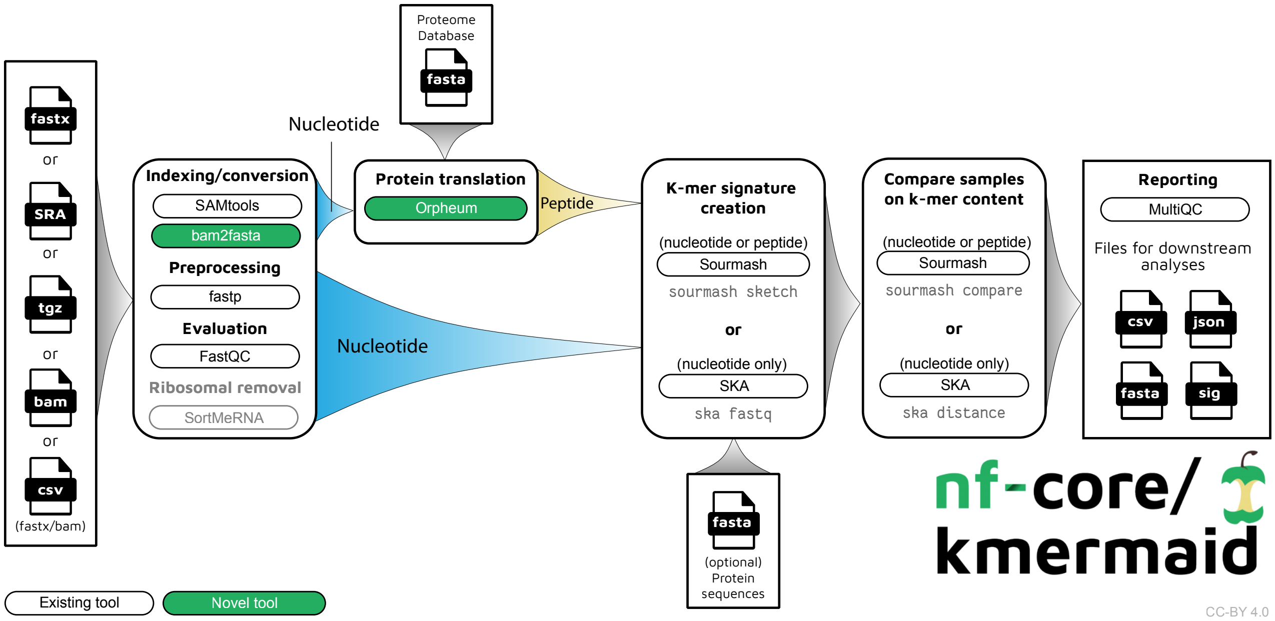


Figure 4: Overview of `nf-core/kmermaid` pipeline to compare DNA/RNA/protein sequences on *k*-mer content. First, any of the following input formats is valid:

1. FASTA or FASTQ (plain text or gzipped)
2. ERA/SRA IDs. The pipeline will automatically retrieve the FASTQ files for the experiment ID using the IDs via nextflow
3. A compressed `.tgz` file containing the exact contents of 10x Genomics' `cellranger count` output, where the `possorted_genome_bam.bam` file is used.
4. An alignment `bam` file
5. A comma-separated variable (`csv`) file containing sample ids and paths to either reads or bams.

If a `.bam` or `.tgz` file are provided, each cell's aligned and unaligned reads are extracted separately. Next, the reads are trimmed, with optional ribosomal removal. If a proteome database in the form of a FASTA file (plain text or compressed as `.gz` is fine) is provided, then translation of the reads with `orpheum` is performed. Next, the *k*-mer signatures are created with Sourmash or SKA. Finally, *k*-mer signatures are optionally compared in an all-by-all computation, creating an $n \times n$ matrix of *k*-mer similarity scores.

Once we have fastq sequence files, performing quality control and trimming using FastQC,[15] read trimming using fastp,[16] optional ribosomal sequence removal with SortMeRNA.[17] Then, nucleotide sequences are translated using Orpheum and a proteome database.

[15] *Babraham Bioinformatics - FastQC A Quality Control tool for High Throughput Sequence Data* n.d.

[16] Chen et al. 2018.

[17] Kopylova, Noé, Pericard, et al. 2014; Kopylova n.d.; Kopylova, Noé, and Touzet 2012.

Both nucleotide and peptide signatures can be created with Sourmash.[18] Split k-mer signatures are possible with Split K-mer Analysis or SKA[19] however, SKA is only applicable to nucleotide *k*-mers and is limited to split *k*-sizes of 9 and 15, or total *k*-mer sizes of 19 and 31. Finally, the samples are compared on their *k*-mer content, creating an $n \times n$ *k*-mer similarity matrix.

[18] Pierce et al. 2019; Brown et al. 2016.

[19] Harris 2018.

#### *Comment on `kmermaid` usage on organisms without reference genomes*

While `kmermaid` uses `.bam` file as input for keeping track of aligned and unaligned *k*-mers, that does *not* mean that one's organism of interest needs to have a reference genome. The main issue is the cell barcode demultiplexing performed by the single-cell RNA-seq processing pipeline, as `kmermaid` creates signatures on exactly the same cellular barcodes as are found in the counts matrix. To confidently assign a single cell barcode to each read, the cell barcode demultiplexing is typically performed on the aligned reads, not the unaligned sequences, and thus for single-cell input to `kmermaid`, it must be in the form of a `.bam` alignment file to separate out individual cell's sequences. The organism's data could be aligned to the any genome to produce aligned/unaligned pairs of sequences, but it may be better to use a closely related species to the organism of interest. This also helps to distinguish between *k*-mers contained in reads that are aligned vs not aligned.

### Sequence data processing

#### Datasets

For mouse, we used droplet lung data from the 18- and 21-month-old mice from *Tabula Muris Senis*.[20] For human, we used droplet lung data from normal lung tissue from the Human Lung Cell Atlas.[21] For bat, we used droplet lung data from both bats from the Chinese horseshoe bat atlas.[22] As the human data has data sharing restrictions due to patient privacy as defined by HIPPA, and the bat data is yet unpublished, at this time, we are only able to share the mouse sequence and *k*-mer signature datasets, available at . Thank you so much for understanding!

[20] Tabula Muris Consortium 2020.

[21] Travaglini et al. 2020.

[22] Ren et al. 2020.

### Cell type database creation

To create cell type databases, we used `sourmash` to merge signatures from single cells labeled as the same cell type, into one cell type signature. To avoid biases due to the numbers of cells per cell type, "noisy" *k*-mers, or *k*-mers that were not shared among at least 5% of cells of that cell type were, removed. To retain only cell type-specific sequences, *k*-mers that were shared by 80% or more cell types were removed. Cell type databases were created using `sourmash` to index sequence bloom trees.

#### Removal of *k*-mers uninformative for cell type annotations

To remove *k*-mers likely uninformative for cell type annotation, we removed sequences from constitutively expressed or single-cell dissociation genes. First, we used the translated protein sequences from GENCODE human v30 and mouse vM21. To cast a wider net, we also used the permissive RefSeq[23] data depository, which contains more sequences than UniProt. RefSeq sequences for Vertebrate Mammalian were downloaded via `rsync` from `http://ftp.ncbi.nlm.nih.gov/refseq/release/vertebrate_mammalian`. Release 98, accessed on 2021-03-08 were used.

[23] O'Leary et al. 2016.

```
rsync \
    --prune-empty-dirs \
    --archive \
    --verbose \
    --recursive \
    --include '*protein.faa.gz' \
    --exclude '/*' \
    rsync://ftp.ncbi.nlm.nih.gov/refseq/release/vertebrate_mammalian/ .
wget https://ftp.ncbi.nlm.nih.gov/refseq/release/RELEASE_NUMBER
DATE=\$(date +'%Y-%M-%d')
RELEASE_NUMBER=\$(cat RELEASE_NUMBER)
zcat *.protein.faa.gz \
```

```
    | gzip -c - \
    > refseq_${RELEASE_NUMBER}_${DATE}_vertebrate_mammalian_concatenated.faa.gz
```

Genes thought to be constitutively expressed were removed using the following regular expression:[24]

[24] Aho 1991.

```
ribosom|mito|ubiqui|ferritin|cytochrome|eukaryotic translation|heat shock|NADH|NADPH
```

The final filtered protein fasta,

```
refseq\_release-98\_\_2020-02-06\_\_vertebrate\_mammalian\_concatenated\_\_nuisance\_genes.fasta.gz
```

is available at `https://osf.io/hgvm5/` (click me to directly download)

To remove nucleotide *k*-mers from constitutive genes, *k*-mers from genes matching the above regular expression from GENCODE human v30 and mouse vM1 transcripts were used, available at `https://osf.io/tjxpn/`.

#### *Merging of single cell signatures and removal of "noisy" k-mers*

To strengthen the cell type signal, single-cell k-mer signatures were merged, first k-mers shared by fewer than 5% of cells per cell type label were removed, then merged into one k-mer signature.

#### *Removal of non cell type-specific k-mers*

Non cell type-specific k-mers shared by 80% or more of the cell types were removed (Supplemental Fig. 2B).

#### *Mouse to mouse database creation and cell type testing*

To prevent overfitting, we created cell type signatures from one male mouse, `MACA_18m_M_LUNG_52` and queried with cells from a female mouse, `MACA_18m_F_LUNG_51`

#### *Mouse cell type database creation for cross-species*

To create a mouse cell type database, we merged cell type signatures from the following mouse IDs from *Tabula Muris Senis*:[25]

[25] Tabula Muris Consortium 2020.

```
MACA_18m_F_LUNG_50
MACA_18m_F_LUNG_51
MACA_18m_M_LUNG_52
MACA_18m_M_LUNG_53
MACA_21m_F_LUNG_54
MACA_21m_F_LUNG_55
```

### References


Aho, Alfred V (1991). *Algorithms for finding patterns in strings, Handbook of theoretical computer science (vol. A): algorithms and complexity.*

Altenhoff, Adrian M, Javier Garrayo-Ventas, Salvatore Cosentino, David Emms, Natasha M Glover, Ana Hernández-Plaza, Yannis Nevers, Vicky Sundesha, Damian Szklarczyk, José M Fernández, Laia Codó, The Quest For Orthologs Consortium, Josep Ll Gelpi, Jaime Huerta-Cepas, Wataru Iwasaki, Steven Kelly, Odile Lecompte, Matthieu Muffato, Maria J Martin, Salvador Capella-Gutierrez, Paul D Thomas, Erik Sonnhammer, and Christophe Dessimoz (2020). "The Quest for Orthologs benchmark service and consensus calls in 2020". en. In: *Nucleic Acids Res.* 48.W1, W538–W545.

*Babraham Bioinformatics - FastQC A Quality Control tool for High Throughput Sequence Data*. `https://www.bioinformatics.babraham.ac.uk/projects/fastqc/`. Accessed: 2021-7-9.

*bam2fasta*.

Boutet, Emmanuel, Damien Lieberherr, Michael Tognolli, Michel Schneider, Parit Bansal, Alan J Bridge, Sylvain Poux, Lydie Bougueleret, and Ioannis Xenarios (2016). "UniProtKB/Swiss-Prot, the Manually Annotated Section of the UniProt KnowledgeBase: How to Use the Entry View". en. In: *Methods Mol. Biol.* 1374, pp. 23–54.

Brown, C Titus and Luiz Irber (2016). "sourmash: a library for MinHash sketching of DNA". In: *JOSS* 1.5, p. 27.

Capella-Gutierrez, Salvador, Adrian Altenhoff, Jaime Huerta-Cepas, Matthieu Muffato, Mateus Patricio, Klaas Vandepoele, Judith Blake, Jesualdo Tomás Fernández Breis, The Quest for Orthologs Consortium, Toni Gabaldón, Erik Sonnhammer, Suzanna Lewis, Quest for Orthologs consortium, Adrian Altenhoff, Carla Bello, Judith Blake, Brigitte Boeckmann, Sébastien Briois, Edward Chalstrey, Hirokazu Chiba, Oscar Conchillo-Solé, Vincent Daubin, Todd DeLuca, Christophe Dessimoz, Jean-Francois Dufayard, Dannie Durand, Ingo Ebersberger, Jesualdo Tomás Fernández-Breis, Kristoffer Forslund, Natasha Glover, Alexander Hauser, Davide Heller, Mateusz Kaduk, Jan Koch, Eugene V Koonin, Evgenia Kriventseva, Shigehiro Kuraku, Odile Lecompte, Olivier Lespinet, Jeremy Levy, Suzanna Lewis, Benjamin Liebeskind, Benjamin Linard, Marina Marcet-Houben, Maria Martin, Claire McWhite, Sergei Mekhedov, Sebastien Moretti, Steven Müller, El-Mabrouk Nadia, Cedric Notredame, Simon Penel, Cécile Pereira, Ivana Pilizota, Henning Redestig, Marc Robinson-Rechavi, Fabian Schreiber, Kimmen Sjölander, Nives Škunca, Alan Sousa da Silva, Martin Steinegger, Damian Szklarczyk, Paul Thomas, Ernst Thuer, Clé-

ment Train, Ikuo Uchiyama, Klaas Vandepoele, Lucas Wittwer, Ioannis Xenarios, Bethan Yates, Evgeny Zdobnov, and Robert M Waterhouse (2017). "Gearing up to handle the mosaic nature of life in the quest for orthologs". In: *Bioinformatics* 34.2, pp. 323–329.
Chen, Shifu, Yanqing Zhou, Yaru Chen, and Jia Gu (2018). "fastp: an ultra-fast all-in-one FASTQ preprocessor". en. In: *Bioinformatics* 34.17, pp. i884–i890.
Consortium, The UniProt (2020). "UniProt: the universal protein knowledgebase in 2021". In: *Nucleic Acids Research* 49.D1, pp. D480–D489. issn: 0305-1048. doi: 10.1093/nar/gkaa1100. eprint: https://academic.oup.com/nar/article-pdf/49/D1/D480/35364103/gkaa1100.pdf. url: https://doi.org/10.1093/nar/gkaa1100.
Crusoe, Michael R., Hussien F. Alameldin, Sherine Awad, Elmar Bucher, Adam Caldwell, Reed Cartwright, Amanda Charbonneau, Bede Constantinides, Greg Edvenson, Scott Fay, Jacob Fenton, Thomas Fenzl, Jordan Fish, Leonor Garcia-Gutierrez, Phillip Garland, Jonathan Gluck, Ivfffdfffdfffdfffdn Gonzfffdfffdfffdfffdlez, Sarah Guermond, Jiarong Guo, Aditi Gupta, Joshua R. Herr, Adina Howe, Alex Hyer, Andreas Hfffdfffdfffdfffdrpfer, Luiz Irber, Rhys Kidd, David Lin, Justin Lippi, Tamer Mansour, Pamela McA'Nulty, Eric McDonald, Jessica Mizzi, Kevin D. Murray, Joshua R. Nahum, Kaben Nanlohy, Alexander Johan Nederbragt, Humberto Ortiz-Zuazaga, Jeramia Ory, Jason Pell, Charles Pepe-Ranney, Zachary N Russ, Erich Schwarz, Camille Scott, Josiah Seaman, Scott Sievert, Jared Simpson, Connor T. Skennerton, James Spencer, Ramakrishnan Srinivasan, Daniel Standage, James A. Stapleton, Joe Stein, Susan R Steinman, Benjamin Taylor, Will Trimble, Heather L. Wiencko, Michael Wright, Brian Wyss, Qingpeng Zhang, en zyme en, and C. Titus Brown (2015). "The khmer software package: enabling efficient nucleotide sequence analysis". In: url: http://dx.doi.org/10.12688/f1000research.6924.1.
Danecek, Petr, James K Bonfield, Jennifer Liddle, John Marshall, Valeriu Ohan, Martin O Pollard, Andrew Whitwham, Thomas Keane, Shane A McCarthy, Robert M Davies, and Heng Li (2021). "Twelve years of SAMtools and BCFtools". en. In: *Gigascience* 10.2.
Dayhoff, Margaret O (1969). *Atlas of protein sequence and structure*. Vol. 4. National Biomedical Research Foundation.
Edgar, Robert C (2004). "Local homology recognition and distance measures in linear time using compressed amino acid alphabets". en. In: *Nucleic Acids Res.* 32.1, pp. 380–385.
Glover, Natasha, Christophe Dessimoz, Ingo Ebersberger, Sofia K Forslund, Toni Gabaldón, Jaime Huerta-Cepas, Maria-Jesus Martin, Matthieu Muffato, Mateus Patricio, Cécile Pereira, Alan Sousa da Silva, Yan Wang, Erik Sonnhammer, and Paul D Thomas (2019).

"Advances and Applications in the Quest for Orthologs". In: *Mol. Biol. Evol.* 36.10, pp. 2157–2164.

Harris, S R (2018). "SKA: Split Kmer Analysis Toolkit for Bacterial Genomic Epidemiology". In: 37.11, pp. 241–224.

Hu, Xiao and Iddo Friedberg (2019). "SwiftOrtho: A fast, memory-efficient, multiple genome orthology classifier". In: *Gigascience* 8.10, pp. 309–312.

Kopylova, Evguenia. *SortMeRNA User Manual*. `http://bioinfo.lifl.fr/RNA/sortmerna/code/SortMeRNA-user-manual.pdf`. Accessed: 2021-7-9.

Kopylova, Evguenia, Laurent Noé, Pierre Pericard, Mikael Salson, and Hélène Touzet (2014). "Sortmerna 2: ribosomal rna classification for taxonomic assignation". In: *Workshop on recent computational advances in metagenomics, ECCB 2014*.

Kopylova, Evguenia, Laurent Noé, and Hélène Touzet (2012). "SortMeRNA: fast and accurate filtering of ribosomal RNAs in metatranscriptomic data". en. In: *Bioinformatics* 28.24, pp. 3211–3217.

Landès, C and J L Risler (1994). "Fast databank searching with a reduced amino-acid alphabet". en. In: *Comput. Appl. Biosci.* 10.4, pp. 453–454.

Leskovec, Jure, Anand Rajaraman, and Jeffrey David Ullman (2020). *Mining of Massive Datasets*. Cambridge, England: Cambridge University Press.

Li, Heng, Bob Handsaker, Alec Wysoker, Tim Fennell, Jue Ruan, Nils Homer, Gabor Marth, Goncalo Abecasis, Richard Durbin, and 1000 Genome Project Data Processing Subgroup (2009). "The Sequence Alignment/Map format and SAMtools". en. In: *Bioinformatics* 25.16, pp. 2078–2079.

Louis, Ard A (2016). "Contingency, convergence and hyper-astronomical numbers in biological evolution". en. In: *Stud. Hist. Philos. Biol. Biomed. Sci.* 58, pp. 107–116.

Murphy, L R, A Wallqvist, and R M Levy (2000). "Simplified amino acid alphabets for protein fold recognition and implications for folding". en. In: *Protein Eng.* 13.3, pp. 149–152.

O'Leary, Nuala A, Mathew W Wright, J Rodney Brister, Stacy Ciufo, Diana Haddad, Rich McVeigh, Bhanu Rajput, Barbara Robbertse, Brian Smith-White, Danso Ako-Adjei, Alexander Astashyn, Azat Badretdin, Yiming Bao, Olga Blinkova, Vyacheslav Brover, Vyacheslav Chetvernin, Jinna Choi, Eric Cox, Olga Ermolaeva, Catherine M Farrell, Tamara Goldfarb, Tripti Gupta, Daniel Haft, Eneida Hatcher, Wratko Hlavina, Vinita S Joardar, Vamsi K Kodali, Wenjun Li, Donna Maglott, Patrick Masterson, Kelly M McGarvey, Michael R Murphy, Kathleen O'Neill, Shashikant Pujar, Sanjida H Rangwala, Daniel Rausch, Lillian D Riddick, Conrad Schoch,

Andrei Shkeda, Susan S Storz, Hanzhen Sun, Francoise Thibaud-Nissen, Igor Tolstoy, Raymond E Tully, Anjana R Vatsan, Craig Wallin, David Webb, Wendy Wu, Melissa J Landrum, Avi Kimchi, Tatiana Tatusova, Michael DiCuccio, Paul Kitts, Terence D Murphy, and Kim D Pruitt (2016). "Reference sequence (RefSeq) database at NCBI: current status, taxonomic expansion, and functional annotation". en. In: *Nucleic Acids Res.* 44.D1, pp. D733–45.

Peterson, Eric L, Jané Kondev, Julie A Theriot, and Rob Phillips (2009). "Reduced amino acid alphabets exhibit an improved sensitivity and selectivity in fold assignment". en. In: *Bioinformatics* 25.11, pp. 1356–1362.

Pierce, N Tessa, Luiz Irber, Taylor Reiter, Phillip Brooks, and C Titus Brown (2019). "Large-scale sequence comparisons with sourmash". en. In: *F1000Res.* 8, p. 1006.

Ren, Lili, Chao Wu, Li Guo, Jiacheng Yao, Conghui Wang, Yan Xiao, Angela Oliveira Pisco, Zhiqiang Wu, Xiaobo Lei, Yiwei Liu, Leisheng Shi, Lianlian Han, Hu Zhang, Xia Xiao, Jingchuan Zhong, Hongping Wu, Mingkun Li, Stephen R Quake, Yanyi Huang, Jianbin Wang, and Jianwei Wang (2020). "Single-cell transcriptional atlas of the Chinese horseshoe bat ( Rhinolophus sinicus) provides insight into the cellular mechanisms which enable bats to be viral reservoirs". In: 6, pp. 23–62.

Simão, Felipe A, Robert M Waterhouse, Panagiotis Ioannidis, Evgenia V Kriventseva, and Evgeny M Zdobnov (2015). "BUSCO: assessing genome assembly and annotation completeness with single-copy orthologs". en. In: *Bioinformatics* 31.19, pp. 3210–3212.

Sonnhammer, Erik L L, Toni Gabaldón, Alan W Sousa da Silva, Maria Martin, Marc Robinson-Rechavi, Brigitte Boeckmann, Paul D Thomas, Christophe Dessimoz, and Quest for Orthologs consortium (2014). "Big data and other challenges in the quest for orthologs". en. In: *Bioinformatics* 30.21, pp. 2993–2998.

Tabula Muris Consortium (2020). "A single-cell transcriptomic atlas characterizes ageing tissues in the mouse". en. In: *Nature* 583.7817, pp. 590–595.

Travaglini, Kyle J, Ahmad N Nabhan, Lolita Penland, Rahul Sinha, Astrid Gillich, Rene V Sit, Stephen Chang, Stephanie D Conley, Yasuo Mori, Jun Seita, Gerald J Berry, Joseph B Shrager, Ross J Metzger, Christin S Kuo, Norma Neff, Irving L Weissman, Stephen R Quake, and Mark A Krasnow (2020). "A molecular cell atlas of the human lung from single-cell RNA sequencing". en. In: *Nature* 587.7835, pp. 619–625.

Waterhouse, Robert M, Mathieu Seppey, Felipe A Simão, Mosè Manni, Panagiotis Ioannidis, Guennadi Klioutchnikov, Evgenia V Kriventseva, and Evgeny M Zdobnov (2018). "BUSCO Applications from Quality Assessments to Gene Prediction and Phylogenomics". en. In: *Mol. Biol. Evol.* 35.3, pp. 543–548.

Ye, Yuzhen, Jeong-Hyeon Choi, and Haixu Tang (2011). "RAPSearch: a fast protein similarity search tool for short reads". en. In: *BMC Bioinformatics* 12, p. 159.

Zhang, Qingpeng, Jason Pell, Rosangela Canino-Koning, Adina Chuang Howe, and C. Titus Brown (2014). "These Are Not the K-mers You Are Looking For: Efficient Online K-mer Counting Using a Probabilistic Data Structure". In: *PLoS ONE* 9.7, e101271. doi: 10.1371/journal.pone.0101271. url: http://dx.doi.org/10.1371%2Fjournal.pone.0101271.
